# Learning recurrent dynamics in spiking networks

**DOI:** 10.1101/297424

**Authors:** Christopher M. Kim, Carson C. Chow

## Abstract

Spiking activity of neurons engaged in learning and performing a task show complex spatiotemporal dynamics. While the output of recurrent network models can learn to perform various tasks, the possible range of recurrent dynamics that emerge after learning remains unknown. Here we show that modifying the recurrent connectivity with a recursive least squares algorithm provides sufficient flexibility for synaptic and spiking rate dynamics of spiking networks to produce a wide range of spatiotemporal activity. We apply the training method to learn arbitrary firing patterns, stabilize irregular spiking activity of a balanced network, and reproduce the heterogeneous spiking rate patterns of cortical neurons engaged in motor planning and movement. We identify sufficient conditions for successful learning, characterize two types of learning errors, and assess the network capacity. Our findings show that synaptically-coupled recurrent spiking networks possess a vast computational capability that can support the diverse activity patterns in the brain.

## Introduction

Neuronal populations exhibit diverse patterns of recurrent activity that can be highly irregular or well-structured when learning or performing a behavioral task^1^–^5^. An open question is whether learning-induced synaptic rewiring is sufficient to give rise to the observed wide range of recurrent spiking dynamics that encodes and processes information.

It has been shown that a network of recurrently connected neuron models can be trained to perform complex motor and cognitive tasks. In this approach, synaptic connections to the outputs are rewired to generate a desired population-averaged signal, while the activity of individual neurons emerges in a self-organized way. The recurrent dynamics resulting from such learning schemes have harnessed chaotic temporally irregular activity of a network of rate-based neurons^6^ that is made repeatable either through direct feedback from the outputs^7,8^ or through training of the recurrent connections directly^9^. The resulting irregular yet stable dynamics provides a rich reservoir from which complex motor commands can be extracted by output neurons that sum over the network linearly^10,11^.

Extending this idea to spiking networks poses a challenge because it is difficult to coordinate the spiking dynamics of many neurons, especially, if spike times are variable as in a balanced network. Some success has been achieved by training spiking networks directly with a feedback loop^12^ or using a rate-based network as an intermediate step^13,14^. A different top-down approach is to build networks that emit spikes optimally to correct the discrepancy between the actual and desired network outputs^15,16^. This optimal coding strategy in a tightly balanced network can be learned with a local plasticity rule^17^ and is able to generate arbitrary network output^18,19^. Although these studies demonstrate that network outputs can perform universal computations^7^, the possible repertoire of the recurrent activity has not been extensively explored. Various network models have been developed to explain certain important features of recurrent dynamics (e.g. fixed point attractor^20^, synchrony^21–23^, sequential activity^3,4,24^, stable chaos^25^, motor planning^26^, and variable spiking^27–30^). However, we still lack an understanding of the scope of recurrent dynamics that spiking networks are capable of generating.

Here we show that a network of spiking neurons is capable of supporting arbitrarily complex coarse-grained recurrent dynamics provided the spatiotemporal patterns of the recurrent activity are diverse, the synaptic dynamics are fast, and the number of neurons in the network is large. We give a theoretical basis for how a network can learn and show various examples, which include stabilizing strong chaotic rate fluctuations in balanced networks and constructing a recurrent network that reproduces the spiking rate patterns of a large number of cortical neurons involved in motor planning and movement. Our study suggests that individual neurons in a recurrent network have the capability to support near universal dynamics.

## Results

### 1 Spiking networks can learn complex recurrent dynamics

We consider a network of *N* quadratic integrate-and-fire neurons that are recurrently connected with spike-activated synapses weighted by a connectivity matrix *W*. We show below that our results do not depend on the spiking mechanism. We focus on two measures of coarse-grained time-dependent neuron activity: 1) the synaptic drive *u*_*i*_(*t*) to neuron *i* which is given by the *W*-weighted sum of low-pass filtered incoming spike trains, and 2) the time-averaged spiking rate *R*_*i*_(*t*) of neuron *i*. The goal is to find a weight matrix *W* that can autonomously generate desired recurrent target dynamics when the network of spiking neurons connected by *W* is stimulated briefly with an external stimulus (**Fig. 1a**). The target dynamics are defined by a set of functions *f*_1_(*t*), *f*_2_(*t*), …, *f*_*N*_(*t*) on a time interval [0, *T*]. Learning of the recurrent connectivity *W* is considered successful if *u*_*i*_(*t*) or *R*_*i*_(*t*) evoked by the stimulus matches the target functions *f*_*i*_(*t*) over the time interval [0, *T*] for all neurons *i* = 1,2,…, *N*.

**Figure 1:**
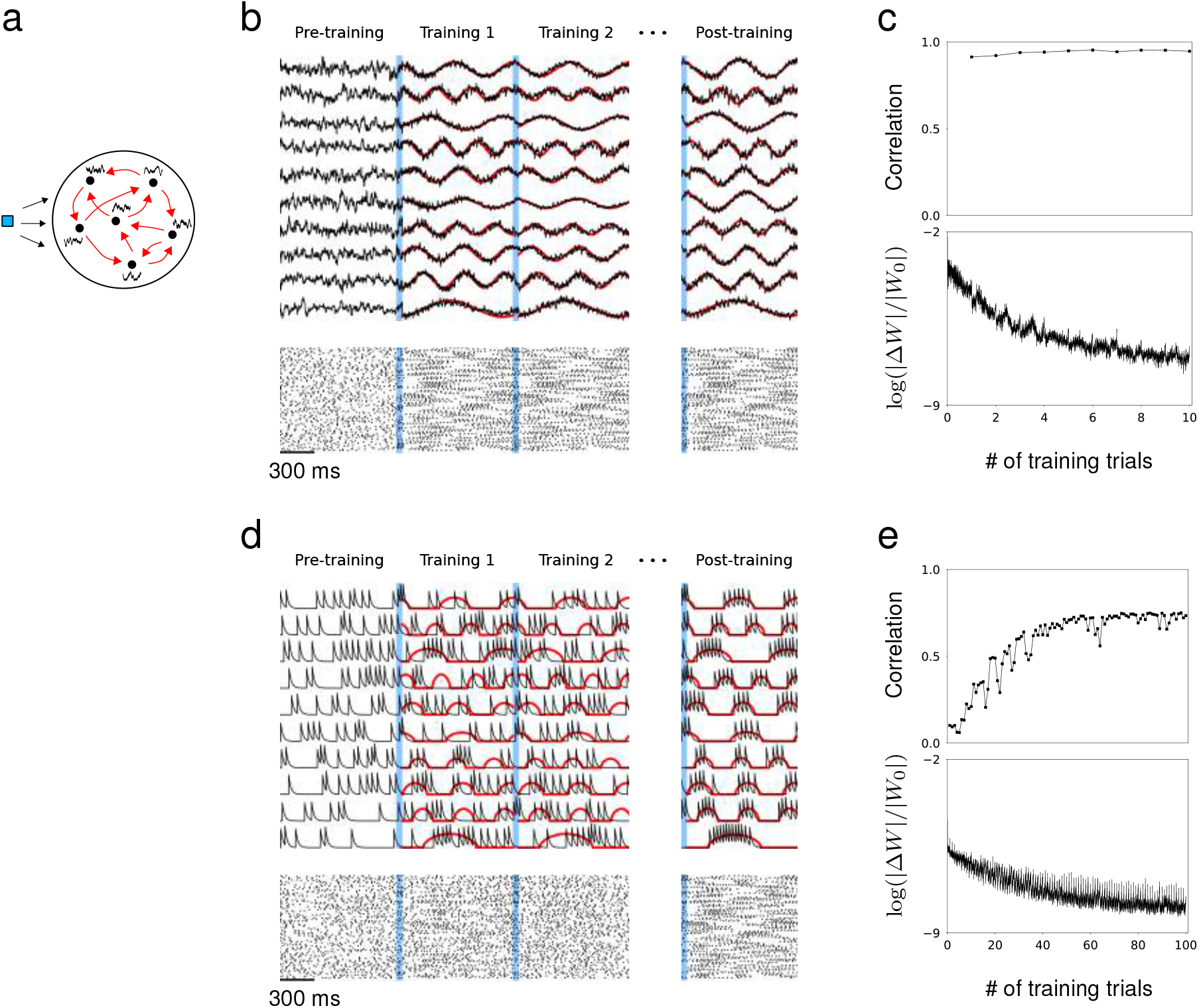
Synaptic drive and spiking rate of neurons in a recurrent network can learn complex patterns. (a) Schematic of network training. Blue square represents the external stimulus that elicits the desired response. Black curves represent target output for each neuron. Red arrows represent recurrent connectivity that is trained to produce desired target patterns. (b) Synaptic drive of 10 sample neurons before, during and after training. Pre-training is followed by multiple training trials. An external stimulus (blue) is applied prior to training for 100 ms. Synaptic drive (black) is trained to follow the target (red). If the training is successful, the same external stimulus can elicit the desired response. Bottom shows the spike rater of 100 neurons. (c) Top, The Pearson correlation between the actual synaptic drive and the target output during training trials. Bottom, The matrix (Frobenius) norm of changes in recurrent connectivity normalized to initial connectivity during training. (d) Filtered spike train of 10 neurons before, during and after training. As in (b), external stimulus (blue) is applied immediately before training trials. Filtered spike train (black) learns to follow the target spiking rate (red) with large errors during the early trials. Applying the stimulus to a successfully trained network elicits the desired spiking rate patterns in every neuron. (e) Top, Same as in (c) but measures the correlation between filtered spike trains and target outputs. Bottom, Same as in (c).

Previous studies have shown that recurrently connected rate units can learn chaotic trajectories that the initial network could already generate^9^, trajectories from a different network^31^, and sequential activity derived from imaging data^32^. Our study expands these results by showing that recurrent dynamics of spiking networks can be trained and the repertoire of recurrent dynamics that can be encoded is vast. To train the recurrent connectivity, we modified the Recursive Least Squares (RLS) algorithm developed in rate models, which minimizes a quadratic cost function between the activity measure and the target together with a quadratic regularization term^8,9,33^ (see Methods).

As a first example, we trained the network to produce synaptic drive patterns that matched a set of sine functions and the spiking rate to match the positive part of the same sine functions. The initial connectivity matrix has connection probability *p* = 0.3 and the coupling strength is drawn from a Normal distribution with mean 0 and standard deviation *σ*. Prior to training, the synaptic drive fluctuates irregularly, but as soon as the RLS algorithm is instantiated, the synaptic drives follow the target with small error; rapid changes in *W* quickly adjust the recurrent dynamics towards the target^8^ (Fig. 1b, c). As a result, the population spike trains exhibit reproducible patterns across training trials. A brief stimulus precedes each training session to reset the network to a specific state. If the training is successful, the trained response can be elicited whenever the same stimulus is applied regardless of the network state. We were able to train a network of rate-based neurons to learn arbitrarily complex target patterns using the same learning scheme (**Supplementary Fig. S1**).

Training the spiking rate is more challenging than training the synaptic drive because small changes in recurrent connectivity do not immediately affect the spiking activity if the effect is below the spike-threshold. Therefore, the spike trains may not follow the desired spiking rate pattern during the early stage of training, and the population spike trains no longer appear similar across training trials (Fig. 1d). This is also reflected in relatively small changes in recurrent connectivity and the substantially larger number of training runs required to produce desired spiking patterns (**Fig. 1e**). However, by only applying the training when the total input to a neuron is suprathreshold, the spiking rate can be trained to reproduce the target patterns. The correlation between the actual filtered spike trains and the target spiking rate increases gradually as the training progresses.

In contrast to training the network read-out, where it is proposed that the initial network needs to be at the “edge of chaos” to learn successfully^8^^12,14,34,35^, the recurrent connectivity can learn to produce the desired recurrent dynamics regardless of the initial network dynamics and connectivity. Even when the initial network has no synaptic connections, the brief stimulus preceding the training session is sufficient to build a fully functioning recurrent connectivity that captures the target dynamics. The RLS algorithm can grow new synapses or tune existing ones as long as some of the neurons become active after the initial stimulus (Supplementary Fig. S2).

Learning was not limited to one set of targets; the same network was able to learn multiple sets of targets. We trained the network to follow two independent sets of targets, where each target function was a sine function with random frequency. Every neuron in the network learned both activity patterns after training, and, when stimulated with the appropriate cue, the network recapitulated the specified trained pattern of recurrent dynamics, regardless of initial activity. The synaptic drive and the spiking rate were both able to learn multiple target patterns (**Fig. 2**).

**Figure 2:**
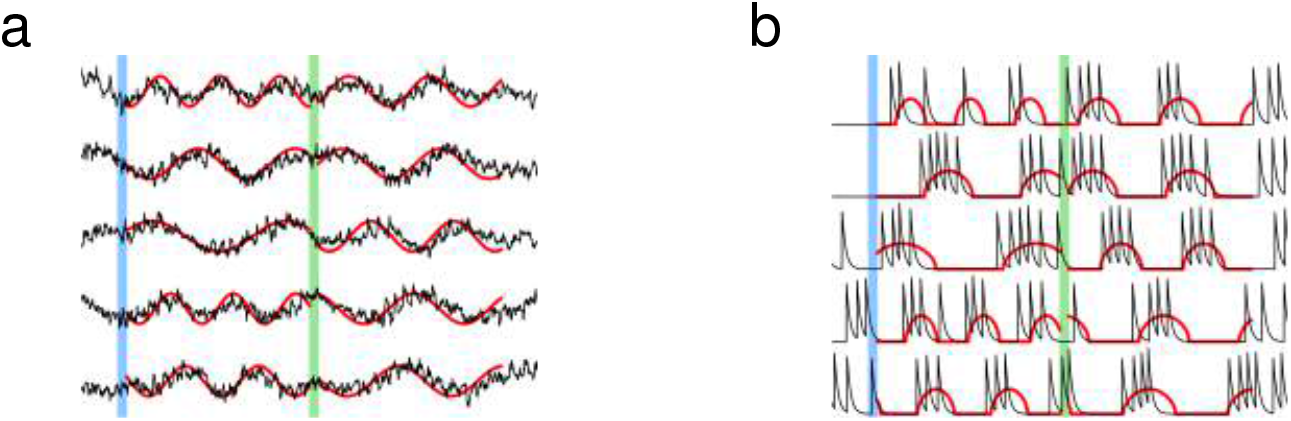
Learning multiple target patterns. (a) The synaptic drive of neurons learns two different target outputs. Blue stimulus evokes the first set of target outputs (red) and the green stimulus evokes the second set of target outputs (red). (b) The spiking rate of individual neurons learns two different target outputs.

### 1.1 Learning arbitrary patterns of activity

To demonstrate that spiking networks can encode recurrent dynamics that have arbitrary spatiotemporal patterns, we considered targets generated from various families of functions; examples include complex periodic functions, chaotic trajectories, and Ornstein-Uhlenbeck (OU) noise. We randomly selected N different target patterns from one of the families to create a set of heterogeneous targets, and trained the synaptic drive of a network consisting of N neurons to learn the target dynamics.

As we will show more rigorously in Section 2, we identified two sufficient conditions on the dynamical state and spatiotemporal structure of target dynamics that ensure a wide repertoire of recurrent dynamics can be learned. The first is a “quasi-static” condition that stipulates that the dynamical time scale of target patterns must be slow enough compared to the synaptic time scale and average spiking rate. The second is a “heterogeneity” condition that requires the spatiotemporal structure of target patterns to be diverse enough. We will quantify these conditions below. The target patterns considered in Figure 3 had slow temporal dynamics in comparison to the synaptic time constant (*τ*_*s*_ = 20 ms) and the patterns were selected randomly to promote diverse structure. After training each neuron’s synaptic drive to produce the respective target pattern, the synaptic drive of every neuron in the network followed its target.

**Figure 3:**
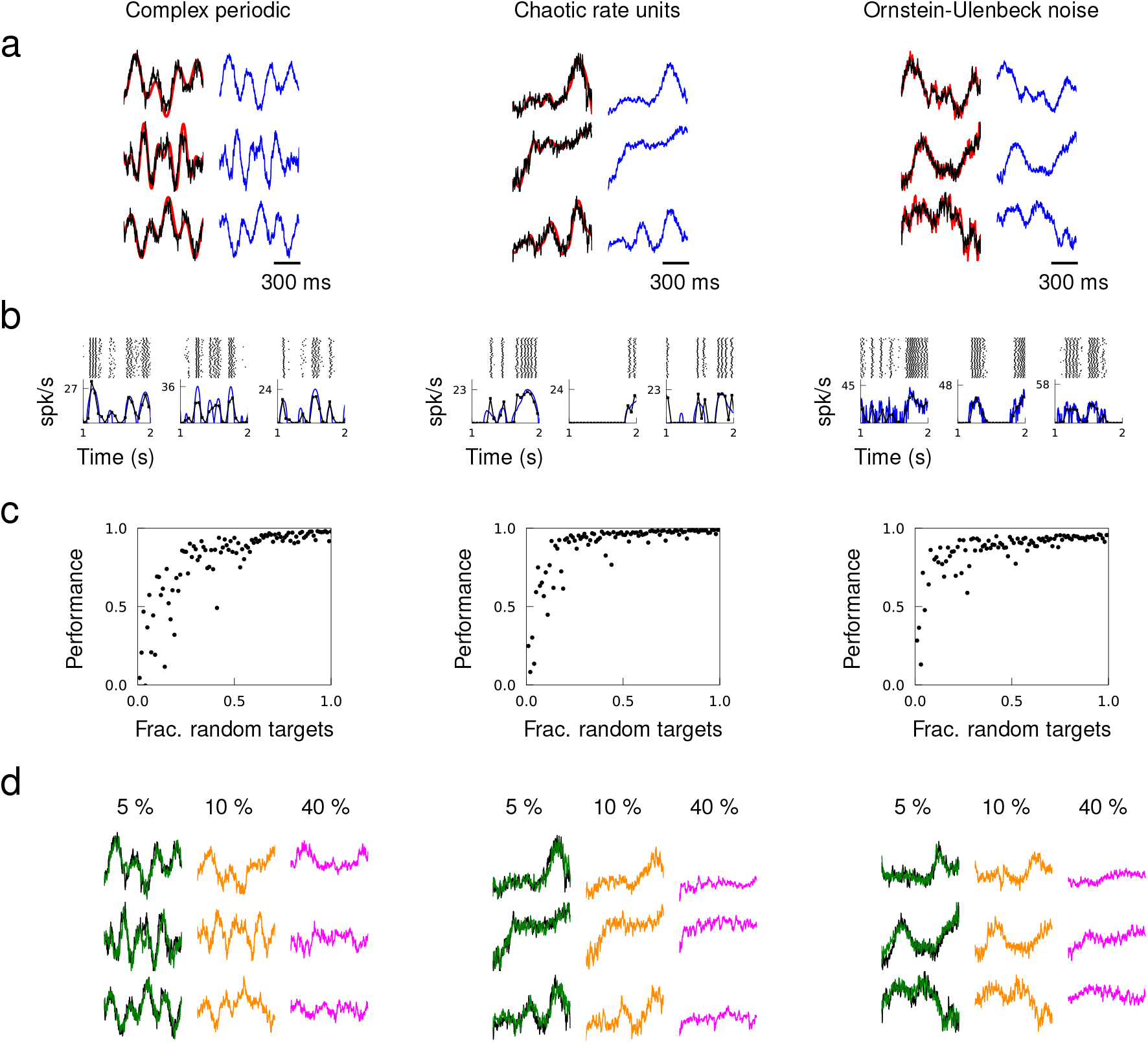
Quasi-static and heterogeneous patterns can be learned. Example target patterns include complex periodic functions (product of sines with random frequencies), chaotic rate units (obtained from a random network of rate units), and OU noise (obtained by low pass filtering white noise with time constant 100 ms). (a) Target patterns (red) overlaid with actual synaptic drive (black) of a trained network. Quasi-static prediction (equation (1)) of synaptic drive (blue). (b) Spike trains of trained neurons elicited multiple trials, trial-averaged spiking rate calculated by the average number of spikes in 50 ms time bins (black), and predicted spiking rate (blue). (c) Performance of trained network as a function of the fraction of randomly selected targets. (d) Network response from a trained network after removing all the synaptic connections from 5%, 10% and 40% of randomly selected neurons in the network.

To verify the quasi-static condition, we compared the actual and a quasi-static approximation of the spiking rate and synaptic drive. The spiking rates of neurons were approximated using the current-to-rate transfer function with time-dependent synaptic input, and the synaptic drive was approximated by a weighted sum of the presynaptic neurons’ spiking rates. We elicited the trained patterns over multiple trials starting at random initial conditions to calculate the trial-averaged spiking rates. The quasi-static approximations of the synaptic drive and spiking rate closely matched the actual synaptic drive (**Fig. 3a**) and trial-averaged spiking rates (**Fig. 3b**).

To examine how the heterogeneity of target patterns may facilitate learning, we created sets of target patterns where the fraction of randomly generated targets was varied systematically. For non-random targets, we used the same target pattern repeatedly. Networks trained to learn target patterns with strong heterogeneity showed that a network is able to encode target patterns with high accuracy if there is a large fraction of random targets (**Fig. 3c**). Networks that are trained on too many repeated target patterns failed to learn. Beyond a certain fraction of random patterns, including additional patterns did not improve the performance, suggesting that the set of basis functions was over-complete. We probed the stability of over-complete networks under neuron loss by eliminating all the synaptic connections from a fraction of neurons. A network was first trained to learn target outputs where all the patterns were selected randomly (i.e. fraction of random targets equals 1) to ensure that the target patterns form a set of a redundant basis functions. Then, we elicited the trained patterns after removing a fraction of neurons from the network, which entails eliminating all the synaptic connections from the lost neurons. A trained network with 5% neuron loss was able to generate the trained patterns perfectly, 10% neuron loss resulted in a mild degradation of network response, and trained patterns completely disappeared after 40% neuron loss (**Fig. 3d**).

The target dynamics considered in **Figure 3** had population spiking rates of 9.1 Hz (periodic), 7.2 Hz (chaotic) and 12.1 Hz (OU) within the training window. To examine how population activity may influence learning, we trained networks to learn target patterns whose average amplitude was reduced gradually across target sets. The networks were able to learn when the population spiking rate of the target dynamics was as low as 1.5 Hz. However, the performance deteriorated as the population spiking rate decreased further (**Supplementary Fig. 2**). To demonstrate that learning does not depend on the spiking mechanism, we trained the synaptic drive of spiking networks using different neuron models. A network of leaky integrate-and-fire neurons, as well as a network of Izhikevich neurons whose neuron parameters were tuned to have five different firing patterns, successfully learned complex synaptic drive patterns (**Supplementary Fig. 3**).

### 1.2 Stabilizing rate fluctuations in balanced networks

A random network with balanced excitation and inhibition is a canonical model for a cortical circuit that produces asynchronous single unit activity^6,27–29,36^. The chaotic activity of balanced rate models^6^ has been harnessed to accomplish complex tasks by including a feedback loop^8^, stabilizing chaotic trajectories^9^ or introducing low-rank structure to the connectivity matrix^37^. Balanced spiking networks have been shown to possess similar capabilities^12–14,16,35^, but it is unknown if it is possible to make use of the heterogeneous fluctuations of the spiking rate in the strong coupling regime^36^. Here, we extend the work of Laje and Buonomano^9^ to spiking networks and show that strongly fluctuating single neuron activities can be turned into dynamic attractors by adjusting the recurrent connectivity.

We take a network with randomly connected excitatory and inhibitory neurons that respects Dale’s Law. Prior to training, the synaptic and spiking activity of individual neurons show large variations across trials because small discrepancies in the initial network state lead to rapid divergence of network dynamics. When simulated with two different initial conditions, the synaptic drive to neurons deviates strongly from each other (**Fig. 4a**), and the spiking activity of single neurons is uncorrelated across trials and the trial-averaged spiking rate has little temporal structure (**Fig. 4b**). The balanced network exhibited sensitivity to small perturbation; the microstate of two identically prepared networks diverged rapidly if one spike is deleted from one of the networks (**Fig. 4c**). It has been previously questioned as to whether the chaotic nature of the balanced state could be utilized to perform reliable computations^25,38^.

**Figure 4:**
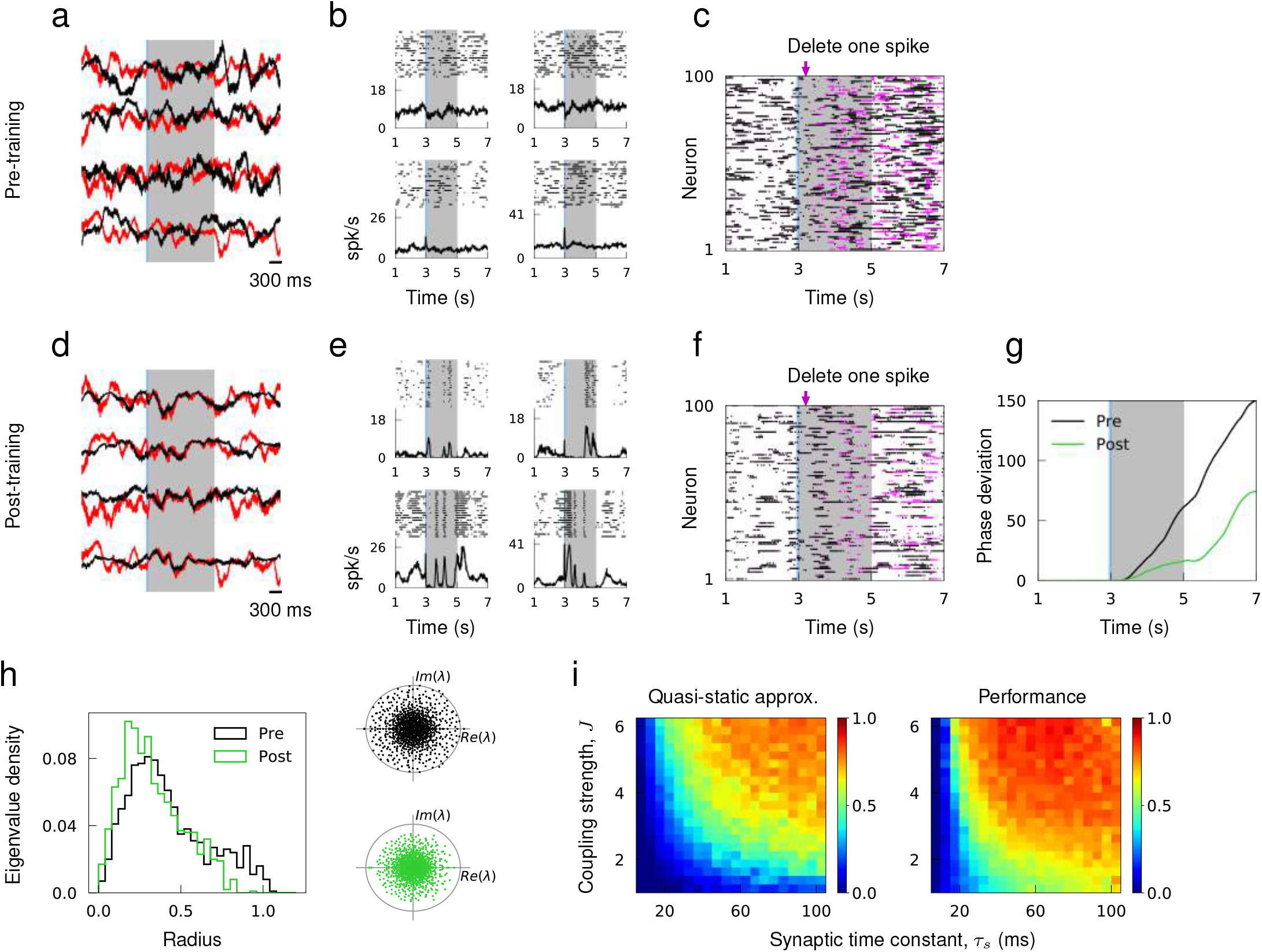
Learning innate activity in balanced networks. (a) Synaptic drive of sample neurons starting at random initial conditions in response to external stimulus prior to training. (b) Spike raster of sample neurons evoked by the same stimulus over multiple trials with random initial conditions. (c) Single spike perturbation of an untrained network. (d)-(f) Synaptic drive, multi-trial spiking response and single spike perturbation in a trained network. (g) The average phase deviation of theta neurons due to single spike perturbation. (h) Left, distribution of eigenvalues of the recurrent connectivity before and after training as a function their absolution values. Right, Eigenvalue spectrum of the recurrent connectivity; gray circle has unit radius. (i) The accuracy of quasi-static approximation in untrained networks and the performance of trained networks as a function of coupling strength J and synaptic time constant *τ*_*s*_. Color bar shows the Pearson correlation between predicted and actual synaptic drive in untrained networks (left) and innate and actual synaptic drive in trained networks (right).

As in Laje and Buonomano^9^, we sought to tame the chaotic trajectories of single neuron activities when the coupling strength is strong enough to induce large and irregular spiking rate fluctuations in time and across neurons^36^. We initiate the untrained network with random initial conditions to harvest innate synaptic activity, i.e. a set of synaptic trajectories that the network already knows how to generate. Then, the recurrent connectivity is trained so that the synaptic drive of every neuron in the network follows the innate pattern when stimulated with an external stimulus. To respect Dale’s Law, the RLS learning rule is modified such that it does not update synaptic connections if there are changes in their signs.

After training, the synaptic drive to every neuron in the network is able to track the innate trajectories in response to the external stimulus within the trained window and diverge from the target pattern outside the trained window (**Fig. 4d**). When the trained network is stimulated to evoke the target patterns, the trial-averaged spiking rate develops a temporal structure that is not present in the untrained network (**Fig. 4e**). To verify the reliability of learned spiking patterns, we simulated the trained network twice with identical initial conditions but deleted one spike 200 ms after evoking the trained response from one of the simulations. Within the trained window, the relative deviation of the microstate is markedly small in comparison to the deviation observed in the untrained network. Outside the trained window, however, two networks diverge rapidly again, which demonstrates that training the recurrent connectivity creates an attracting flux tube around what used to be chaotic spike sequences^25^ (**Fig. 4f, g**). Analyzing the eigenvalue spectrum of the recurrent connectivity reveals that the distribution of eigenvalues shifts towards zero and the spectral radius decreases as a result of training, which is consistent with the more stable network dynamics found in trained networks.

To demonstrate that learning the innate trajectories works well when a balanced network satisfies the quasi-static condition, we scanned the coupling strength *J* (see Figure Methods for definition of *J*) and synaptic time constant *τ*_s_ over a wide range and evaluated the accuracy of the quasi-static approximation in untrained networks. We find that increasing either *J* or *τ*_*s*_ promotes strong fluctuations in spiking rates^36,39^, hence improving the quasi-static approximation (**Fig. 4i**). Learning performance was correlated with adherence to the quasi-static approximation, resulting in better performance for strong coupling and long synaptic time constants.

### 1.3 Generating an ensemble of in-vivo spiking patterns

We next investigated if the training method applies to actual spike recordings of a large number of neurons. In a previous study, a network of rate units was trained to match sequential activity imaged from posterior parietal cortex as a possible mechanism for short-term memory^3,32^. Here, we aimed to construct recurrent spiking networks that capture heterogeneous spiking activity of cortical neurons involved in motor planning and movement^1,2,5^.

The *in-vivo* spiking data was obtained from the publicly available data of Li et al.^5^, where they recorded the spike trains of a large number of neurons from the anterior lateral motor cortex of mice engaged in planning and executing directed licking over multiple trials. We compiled the trial-average spiking rate of *N*_cor_ = 227 cortical neurons from their data set^40^, and trained a recurrent network model to reproduce the spiking rate patterns of all the *N*_cor_ neurons autonomously in response to a brief external stimulus. We only trained the recurrent connectivity and did not alter single neuron dynamics or external inputs.

First, we tested if a recurrent network of size *N*_cor_ is able to generate the spiking rate patterns of the same number of cortical neurons. This network model assumes that the spiking patterns of *N*_cor_ cortical neurons can be self-generated within a recurrent network. After training, the spiking rate of neuron models captured the overall trend of spiking rate, but not the rapid changes that may be pertinent to the short term memory and motor response (**Fig. 5b**). We hypothesized that the discrepancy may be attributed to other sources of input to the neurons not included in the model, such as recurrent input from other neurons in the local population or input from other areas of the brain, or the neuron dynamics that cannot be captured by our neuron model. We thus sought to improve the performance by adding *N*_aux_ auxiliary neurons to the recurrent network to mimic the spiking activity of unobserved neurons in the local population, and trained the recurrent connectivity of a network of size *N*_cor_ + *N*_aux_ (**Fig. 5a**). The auxiliary neurons were trained to follow spiking rate patterns obtained from an OU process and provided heterogeneity to the overall population activity patterns. When *N*_aux_/*N*_cor_ ≥ 2, the spiking patterns of neuron models accurately fit that of cortical neurons (**Fig. 5c**), and the population activity of all *N*_cor_ cortical neurons was well captured by the network model (**Fig. 5d**). The fit to cortical activity improved gradually as a function of the fraction of auxiliary neurons in the network due to increased heterogeneity in the target patterns (**Fig. 5e**)

**Figure 5:**
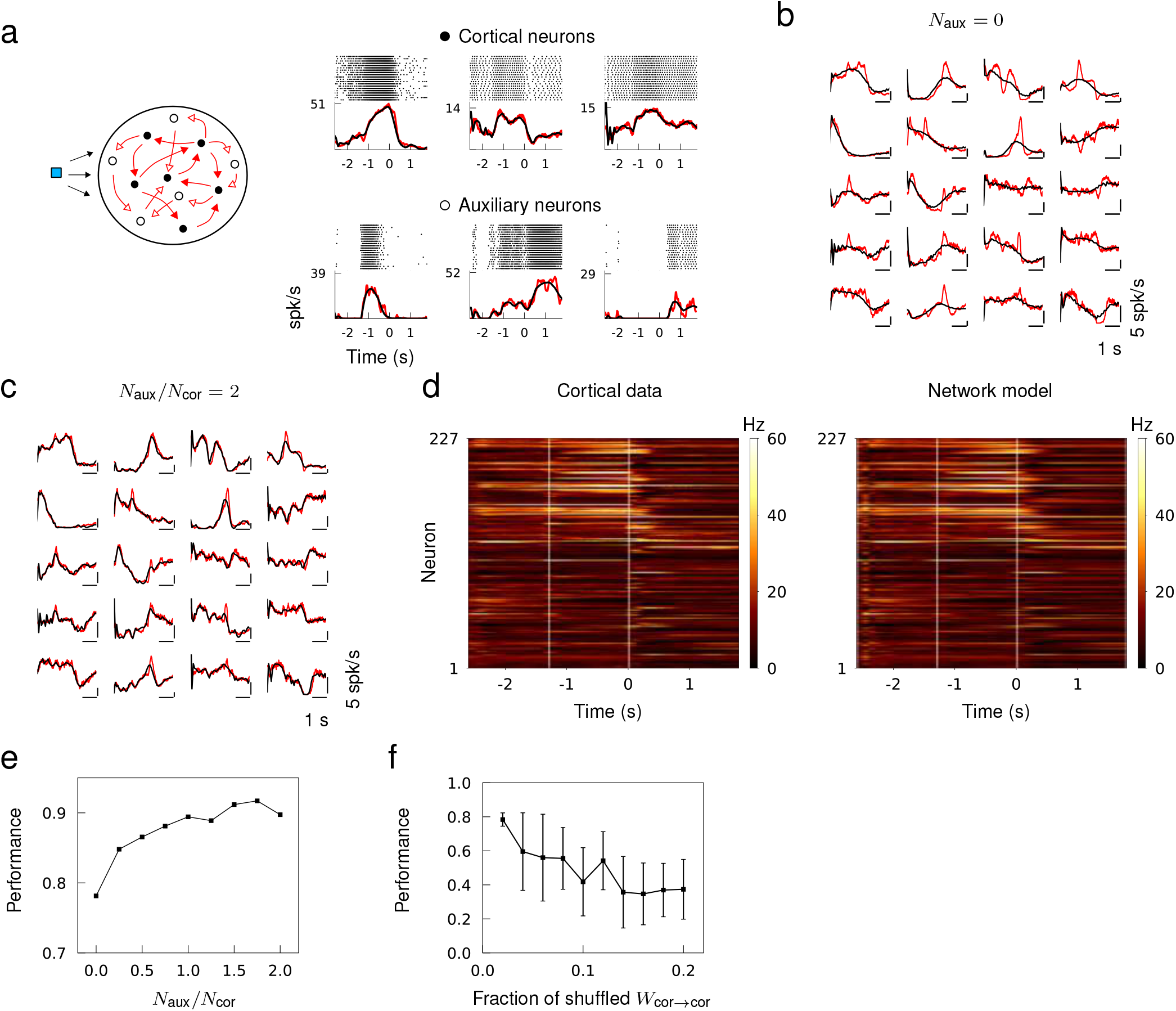
Generating *in-vivo* spiking activity in a subnetwork of a recurrent network. (a) Network schematic showing cortical (black) and auxiliary (white) neuron models trained to follow the spiking rate patterns of cortical neurons and target patterns derived from OU noise, respectively. Multi-trial spike sequences of sample cortical and auxiliary neurons in a successfully trained network. (b) Trial-averaged spiking rate of cortical neurons (red) and neuron models (black) when no auxiliary neurons are included. (c) Trial-averaged spiking rate of cortical and auxiliary neuron models when *N*_aux_/*N*_cor_ = 2. (c) Spiking rate of all the cortical neurons from the data (left) and the recurrent network model (right) trained with *N*_aux_/*N*_cor_ = 2. (e) The fit to cortical dynamics improves as the number of auxiliary neurons increases. (f) Random shuffling of synaptic connections between cortical neuron models degrades the fit to cortical data. Error bars show the standard deviation of results from 10 trials.

To verify that the cortical neurons in the network model are not simply driven by the feedforward inputs from the auxiliary neurons, we randomly shuffled a fraction of recurrent connections between cortical neurons after a successful training. The fit to cortical data deteriorated as the fraction of shuffled synaptic connections between cortical neurons was increased, which confirmed that the recurrent connections between the cortical neurons played a role in generating the spiking patterns (**Fig. 5f**).

## 2 Sufficient conditions for learning

Here we quantify the sufficient conditions the target patterns need to satisfy in order to be successfully encoded in a network. The first condition is that the target patterns must be sufficiently slow compared to the dynamical time scale of both neurons and synapses, such that targets can be considered constant (quasi-static) on a short time interval. As we show below, this is not overly restrictive. In terms of network dynamics, the quasi-static condition implies that the synaptic and neuron dynamics operate as if in a stationary state even though the stationary values change as the network activity evolves in time. In this quasi-static state, we can use a mean field description of the spiking dynamics to derive a self-consistent equation that captures the time-dependent synaptic and spiking activity of neurons^22,36,41^ (see **Supplementary Notes**). Under the quasi-static approximation, the synaptic drive satisfies

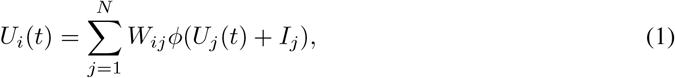

and the spiking rate *R*_*i*_ = *φ*(*U*_*i*_ + *I*_*i*_) satisfies

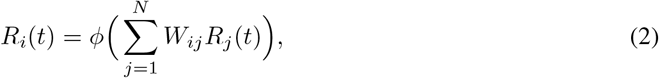

where *φ* is the current-to-rate transfer (i.e. gain) function and *I*_*i*_ is a constant external input.

The advantage of operating in a quasi-static state is that both measures of network activity become conducive to learning new patterns. First, equation (1) is closed in terms of *U,* which implies that training the synaptic drive is equivalent to training a rate-based network. Second, the RLS algorithm can efficiently optimize the recurrent connectivity *W,* thanks to the linearity of equation (1) in *W,* while the synaptic drive closely follows the target patterns as shown in Fig. 1b. The spiking rate also provides a closed description of the network activity, as described in equation (2). However, due to nonlinearity in *W,* it learns only when the total input to a neuron is supra-threshold, i.e. the gradient of *φ* must be positive. For this reason, the learning error cannot be controlled as tightly as the synaptic drive and requires additional trials for successful learning as shown in **Fig. 1d**.

The second condition requires the target patterns to be sufficiently heterogeneous in time and across neurons. Such complexity allows the ensemble of spiking activity to have a rich spatiotemporal structure to generate the desired activity patterns of every neuron within the network. In the perspective of “reservoir computing”^8,11,42^, every neuron in a recurrent network is considered to be a read-out, and, at the same time, part of the reservoir that is collectively used to produce desired patterns in single neurons. The heterogeneity condition is equivalent to having a set of complete (or over-complete) basis functions, i.e. φ(*U*_*j*_ + *I*_*j*_), *j* = 1,…, *N* in equation (1) and *R*_*j*_, *j* = 1,…, *N* in equation (2), to generate the target patterns, i.e. the left hand side of equations (1) and (2). The two conditions are not necessarily independent. Heterogeneous targets also foster asynchronous spiking activity that support quasi-static dynamics.

We note that, although equations (1) and (2) describe the dynamical state in which the learning works well, merely finding *W* that satisfies one of the equations does not guarantee that a spiking network with recurrent connectivity *W* will produce the target dynamics. The recurrent connectivity *W* needs to be trained iteratively as the network dynamics unfold in time to ensure that the target dynamics can be generated in a stable manner^8^. Moreover, the equations explain how the target dynamics can be learned, but does not address how recurrent spiking networks are able to encode such dynamic patterns.

### 2.1 Characterizing learning error

Learning errors can be classified into two categories. There are *tracking* errors, which arise because the target is not a solution of the true spiking network dynamics and *sampling* errors, which arise from encoding a continuous function with a finite number of spikes. We quantified these learning errors as a function of the network and target time scales. The intrinsic time scale of spiking network dynamics is the synaptic decay constant *τ*_s_, and the time scale of target dynamics is the decay constant *τ*_*c*_ of OU noise. We used target patterns generated from OU noise since the trajectories have a predetermined time scale and their spatio-temporal patterns are sufficiently heterogeneous.

We systematically varied *τ*_*s*_ and *τ*_*c*_ from fast AMPA-like (~ 1ms) to slow NMDA-like synaptic transmission (~ 100ms) and trained the synaptic drive of networks with synaptic time scale *τ*_*s*_ to learn a set of OU trajectories with time scale *τ*_*c*_. The parameter scan reveals a learning regime, where the networks successfully encode the target patterns, and two error-dominant regimes. The tracking error is prevalent when synapses are slow in comparison to target patterns, and the sampling error dominates when the synapse is fast (**Fig. 6a**).

A network with a synaptic decay time *τ*_*s*_ = 200 ms fails to follow rapid changes in the target patterns, but still captures the overall shape, when the target patterns have a faster time scale *τ*_*c*_ = 100 ms (Fig. 6b, **Tracking error**). This prototypical example shows that the synaptic dynamics are not fast enough to encode the target dynamics in the tracking error regime. With a faster synapse *τ*_*s*_ = 30 ms, the synaptic drive is able to learn the identical target trajectories with high accuracy (**Fig. 6b, Learning**). However, when the synapse is too fast *τ*_*s*_ = 5 ms, the synaptic drive fluctuates around the target trajectories with high frequency (Fig. 6b, Sampling error). This is a typical network response in the sampling error regime since discrete spikes with narrow width and large amplitude are summed to “sample” the target synaptic activity.

**Figure 6:**
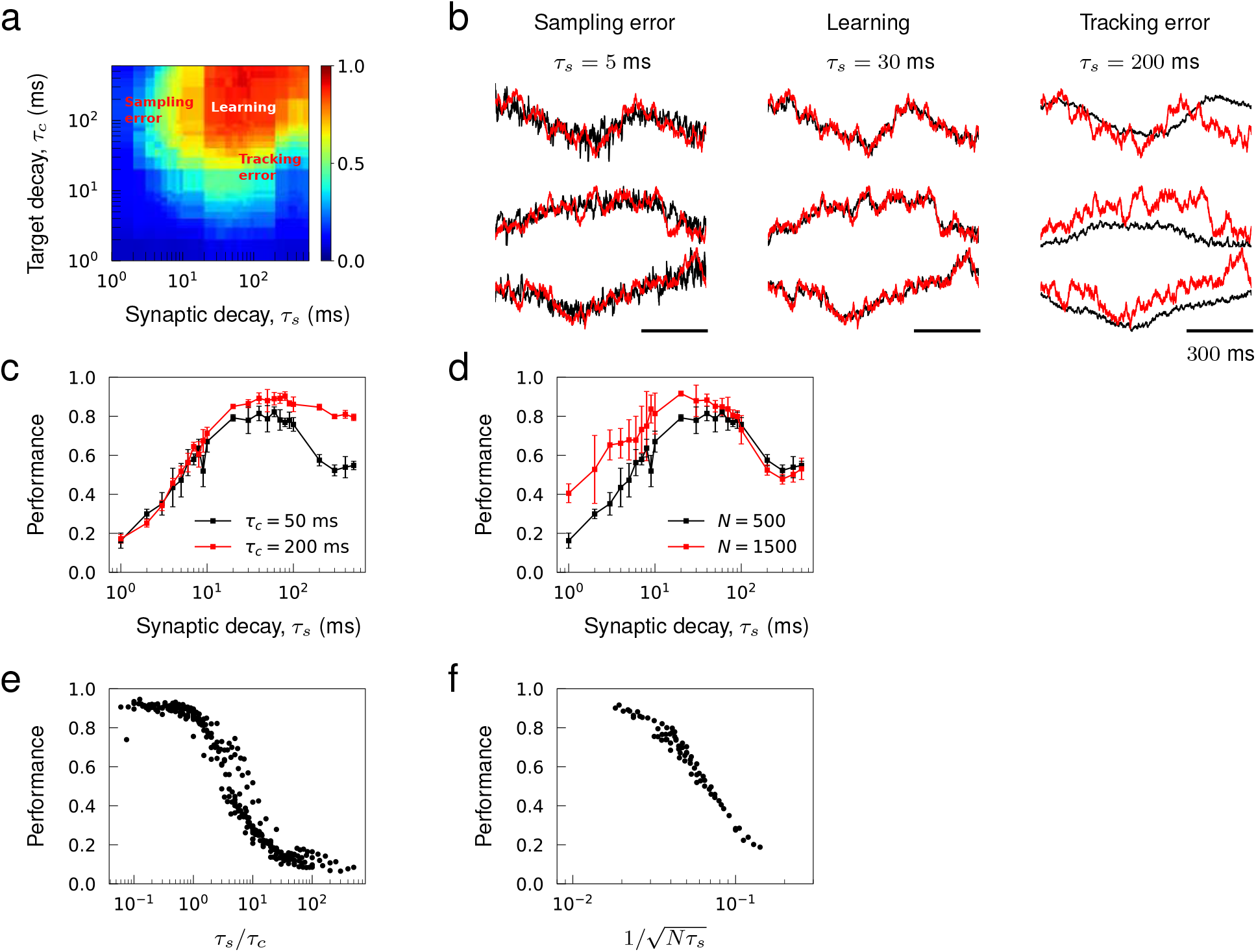
Sampling and tracking errors. Synaptic drive was trained to learn 1 s long trajectories generated from OU noise with decay time *τ*_*c*_. (a) Performance of networks of size *N* = 500 as a function of synaptic decay time *τ*_*s*_ and target decay time *τ*_*c*_. (b) Examples of trained networks whose responses show sampling error, tracking error, and successful learning. The target trajectories are identical and *τ*_*c*_ = 100 ms. (c) Inverted “U”-shaped curve as a function of synaptic decay time. Error bars show the s.d. of five trained networks of size *N* = 500. (d) Inverted “U”-shaped curve for networks of sizes *N* = 500 and 1000 for *τ*_*c*_ = 100 ms. (e) Network performance shown as a function of *τ*_*s*_/*τ*_*c*_ where the range of *τ*_*s*_ is from 30 ms to 500 ms and the range of *τ*_*c*_ is from 1 ms to 500 ms and *N =* 1000. (**f**) Network performance shown as a function of 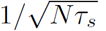 where the range of *τ*_*s*_ is from 1 ms to 30 ms, the range of *N* is from 500 to 1000 and *τ*_*c*_ = 100 ms.

To better understand how network parameters determine the learning errors, we mathematically analyzed the errors assuming that (1) target dynamics can be encoded if the quasi-static condition holds, and (2) the mean field description of the target dynamics is accurate (see Supplementary Notes). The learning errors were characterized as a deviation of these assumptions from the actual spiking network dynamics. We found that the tracking errors ∊_track_ are substantial if the quasi-static condition is not valid, i.e. synapses are not fast enough for spiking networks to encode targets, and the sampling errors ∊_sample_ occur if the mean field description becomes inaccurate, i.e. discrete representation of targets in terms of spikes deviates from their continuous representation in terms of spiking rates. The errors are estimated to scale with

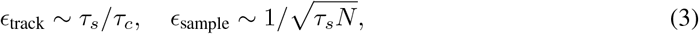

which imply that tracking error can be controlled as long as synapses are relatively faster than target patterns, and the sampling error can be controlled by either increasing *τ*_*s*_ to stretch the width of individual spikes or increasing *N* to encode the targets with more input spikes. The error estimates reveal the versatility of recurrent spiking networks to encode arbitrary patterns since *∊*_track_ can be reduced by tuning *τ*_*s*_ to be small enough and *∊*_sample_ can be reduced by increasing *N* to be large enough.

**Figure 7:**
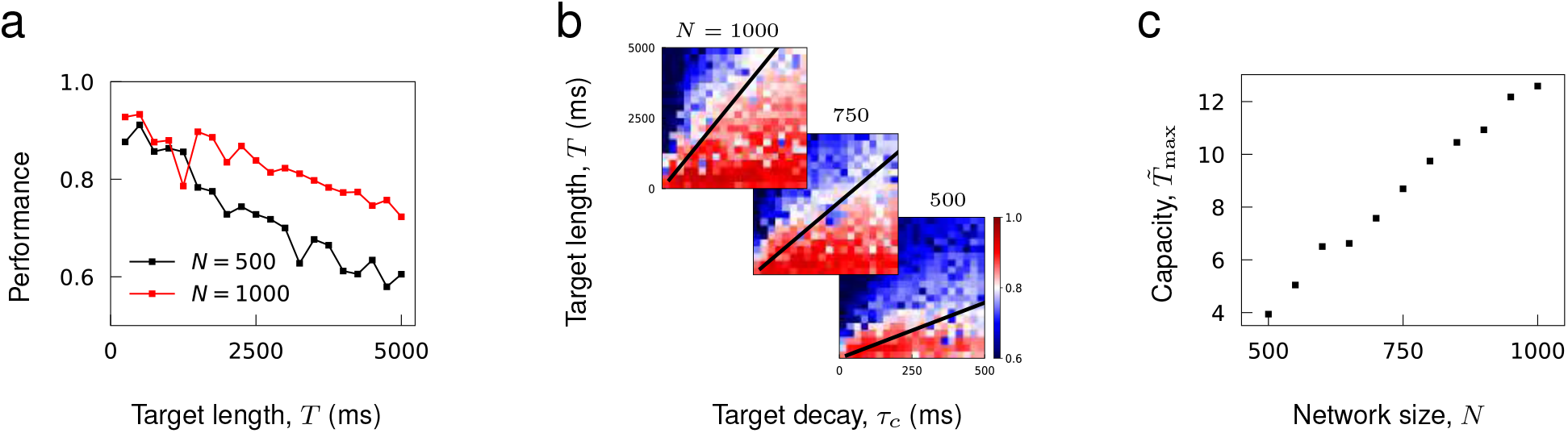
Capacity as a function of network size. (a) Performance of trained networks as a function of target length *T* for networks of size *N =* 500 and 1000. Target patterns were generated from OU noise with decay time *τ*_*c*_ = 100 ms. (**b**) Networks of fixed sizes trained on a range of target length and correlations. Color bar shows the Pearson correlation between target and actual synaptic drive. The black lines show the function 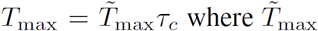 was fitted to minimize the least square error between the linear function and maximal target length *T*_max_ that can be successfully learned at each *τ*_*c*_. (c) Learning capacity 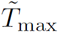 shown as a function of network size.

We examined the performance of trained networks to verify if the theoretical results can explain the learning errors. The learning curve, as a function of *τ*_*s*_, has an inverted U-shape when both types of errors are present (**Fig. 6c, d**). Successful learning occurs in an optimal range of *τ*_*s*_, and, consistent with the error analysis, the performance decreases monotonically with *τ*_*s*_ on the right branch due to the tracking error while increases monotonically with *τ*_*s*_ on the left branch due to the sampling error. The tracking error is reduced if target patterns are slowed down from *τ*_*c*_ = 50 ms to *τ*_*c*_ = 200 ms, hence decrease the ratio *τ*_*s*_/*τ*_*c*_. Then, the learning curve becomes sigmoidal, and the performance remains high even when *τ*_*s*_ is in the slow NMDA regime (Fig. 6c). On the other hand, the sampling error is reduced if the network size is increased from *N* = 500 to 1500, which lifts the left branch of the learning curve (Fig. 6d). Note that when two error regimes are well separated, changes in target time scale *τ*_*c*_ does not affect *∊*_sample_, and changes in network size *N* does not affect *∊*_sample_, as predicted.

Finally, we condensed the training results over a wide range of target time scales in the tracking error regime (**Fig. 6e**), and similarly condensed the training results over different network sizes in the sampling error regime (**Fig. 6f**) to demonstrate that *τ*_*s*_/*τ*_*c*_ and *Nτ*_*s*_ explain the overall performance in the tracking and sampling error regimes, respectively.

### 2.2 Learning capacity increases with network size

It has been shown that a recurrent rate network’s capability to encode target patterns deteriorates as a function of the length of time^9^, but increase in network size can enhance its storage capacity^32,43,44^. Consistent with these results, we find that the performance of recurrent spiking networks to learn complex trajectories decreases with target length and improves with network size (**Fig. 7a**).

To assess the storage capacity of spiking networks, we evaluated the maximal target length that can be encoded in a network as a function of network size. It was necessary to define the target length in terms of its “effective length” to account for the fact that target patterns with the same length may have different effective length due to their temporal structures; for instance, OU noise with short temporal correlation times has more structure to be learned than a constant function. For target trajectories generated from an OU process with decay time *τ*_*c*_, we rescaled the target length *T* with respect to *τ*_*c*_ and defined the effective length 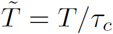. The capacity of a network is the maximal 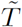 that can be successfully encoded in a network.

To estimate the maximal 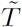, we trained networks of fixed size to learn OU trajectories while varying *T* and *τ*_*c*_ (each panel in **Fig. 7b**). Then, for each *τ*_*c*_, we find the maximal target length *T*_max_ that can be learned successfully, and estimate the maximal 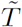 by finding a constant 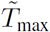 that best fits the line 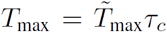 to training results (black lines in **Fig. 7b**). **Figure 7c** shows that the learning capacity 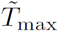 increases monotonically with the network size.

We can illustrate why this may be the case with a simple albeit non-rigorous counting argument. Successful learning is achieved for the synaptic drive when equation (1) is satisfied. If we artificially discretize time into *P* “quasi-static” bins then we can consider *U*_*i*_(*t*) as a *N × P* matrix that satisfies a matrix equation *U = WV,* where we treat *V* ≡ *φ*(*U* + *I*) as an independent matrix. If we operate on the matrix equation with *V*^*T*^ we find that a solution to this artificial system is possible if the matrix *VV*^*T*^ is invertible, which is possible if the rank of *VV*^*T*^ is full. This argument then shows how increasing pattern heterogeneity, which makes the rows of *V* less correlated, is beneficial for learning, and that learning capability will decline as *P* increases, with a steep decline for *P > N*. If we ascribe quasi-static bin to some fraction of the pattern correlation time then *P* will scale with the length of the time interval. In this way, we can intuitively visualize the temporal storage capacity.

## Discussion

We have shown that the synaptic drive and spiking rate of a network of spiking neurons can be trained to follow any arbitrary spatiotemporal patterns. The necessary ingredients for learning are that the spike train inputs to a neuron are weakly correlated (i.e. heterogeneous target patterns), the synapses are fast enough (i.e. small tracking error), and the network is large enough (i.e. small sampling error and large capacity). We demonstrated that (1) a balanced network consisting of excitatory and inhibitory neurons can learn to track its strongly fluctuating innate synaptic trajectories, and (2) a recurrent spiking network can learn to reproduce the spiking rate patterns of an ensemble of cortical neurons involved in motor planning and movement.

Our scheme works because the network quickly enters a quasi-static state where the instantaneous firing rate of a neuron is a fixed function of the inputs. Learning fails if the synaptic time scale is slow compared to the time scale of the target, in which case the quasi-static condition is violated and the tracking error becomes large. There is a trade-off between tracking error and sampling noise; fast synapse can decrease the tracking error, but it also increases the sampling noise. Increasing the network size can decrease sampling noise without affecting the tracking error. Therefore, by adjusting the synaptic time scale and network size, it becomes possible to learn arbitrarily complex recurrent dynamics.

An important structural property of our network model is that the synaptic inputs are summed linearly, which allows the synaptic activity to be trained using a recursive form of linear regression^8^. Linear summation of synaptic inputs is a standard assumption for many spiking network models^21,22,27–29^ and there is physiological evidence that linear summation is prevalent^45,46^. Training the spiking rate, on the other hand, does not take full advantage of the linear synapse due to the nonlinear current-to-transfer function. The network is capable of following a wide repertoire of patterns because even though the network dynamics are highly nonlinear, the system effectively reduces to a linear system for learning. Moreover, learning capacity can be estimated using a simple solvability condition for a linear system. However, nonlinear dendritic processing has been widely observed^47,48^ and may have computational consequences^14,49,50^. It requires further investigation to find out whether a recurrent network with nonlinear synapses can be trained to learn arbitrary recurrent dynamics.

We note that our training scheme does not teach precise spike timings; it either trains the spiking rate directly or trains the synaptic drive which in turn trains the spiking rate. The spike times can be divergent from instance to instance, hence our learning scheme supports rate coding as opposed to spike coding; we demonstrated that the same learning scheme can be used to train the recurrent dynamics of rate-based networks. We find that spike trains that have temporally irregular structure across neurons enhance the rate coding scheme by providing sufficient computational complexity to encode the target dynamics. All neurons in the network can be trained to follow the same synaptic drive patterns as long as there is sufficient heterogeneity, e.g. noisy external input, that decorrelates the spike trains of the neurons (**Supplementary Fig. S5**).

Although our results confirm that recurrent spiking networks have the capability to generate a wide range of repertoire of recurrent dynamics, it is unlikely that a biological network is using this particular learning scheme. The learning rule derived from recursive least squares algorithm is very effective but is nonlocal in time, i.e. it uses the activity of all presynaptic neurons within the train time window to update synaptic weights. Moreover, each neuron in the network is assigned with a target signal and the synaptic connections are updated at a fast time scale as the error function is computed in a supervised manner. It would be of interest to find out whether more biologically plausible learning schemes, such as reward-based learning^51–53^, can lead to similar performance.

Our study provided conditions under which a spiking network can learn a wide range of target dynamics. Previous studies investigated the mathematical relationship between the patterns of stored fixed points and the recurrent connectivity in simple network models^54,55^. It remains an open question to classify possible target patterns that can be encoded in a spiking network and identify the class of connectivity matrices that generates the target patterns in a dynamically stable manner.

## Methods

### 1 Network of spiking neurons

We considered a network of *N* randomly and sparsely connected quadratic integrate-and-fire neurons given by

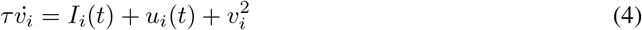

where *v*_*i*_ is a dimensionless variable representing membrane potential, *I*_*i*_(*t*) is an applied input, *u*_*i*_(*t*) is the total synaptic drive the neuron receives from other neurons in the recurrent network, and *τ* = 10 ms is a neuron time constant. The threshold to spiking is zero input. For negative total input, the neuron is at rest and for positive input, *v*_*i*_ will go to infinity or “blow up” in finite time from any initial condition. The neuron is considered to spike at *v*_*i*_ = ∞ whereupon it is reset to – ∞^56,57^.

To simulate the dynamics of quadratic integrate-and-fire neurons, we used its phase representation, i.e. theta neuron model, that can be derived by a simple change of variables, *v*_*i*_ = tan(*θ*_*i*_/2); its dynamics are governed by

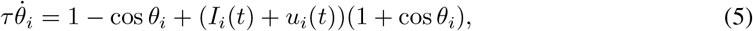

where a spike is emitted when *θ*(*t*) = π. The synaptic drive to a neuron obeys

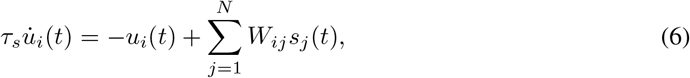

where 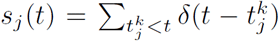 is the spike train neuron *j* generates up to time *t,* and *τ*_*s*_ is a synaptic time constant.

The recurrent connectivity *W*_*ij*_ describes the synaptic coupling from neuron *j* to neuron *i*. It can be any real matrix but in many of the simulations we use a random matrix with connection probability *p,* and the coupling strength of non-zero elements is modeled differently for different figures.

### 2 Training recurrent dynamics

To train the synaptic and spiking rate dynamics of individual neurons, it is more convenient to divide the synaptic drive equation (6) into two parts; one that isolates the spike train of single neuron and computes its synaptic filtering

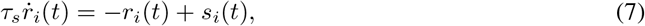

and the other that combines all the presynaptic neurons’ spiking activity and computes the synaptic drive

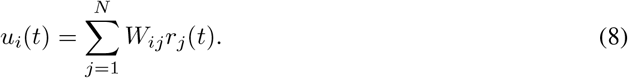

The synaptic drive u_i_ and the filtered spike train *r*_*i*_ are two measures of spiking activity that have been trained in this study. Note that equations (7) and (8) generate synaptic dynamics that are equivalent to equation (6).

#### Training procedure

We select *N* target trajectories *f*_1_(*t*),…, *f*_*N*_(*t*) of length *T* ms for a recurrent network consisting of *N* neurons. We train either the synaptic drive or spiking rate of individual neuron *i* to follow the target *f*_*i*_(*t*) over time interval [0, *T*] for alli = 1, *…,N*. External stimulus *I*_*i*_ with amplitude sampled uniformly from [–1,1] is applied to neuron *i* for all *i* = 1, 2,…, *N* for 100 ms immediately preceding the training to situate the network at a specific state. During training, the recurrent connectivity *W* is updated every ∆*t* ms using a learning rule described below in order to steer the network dynamics towards the target dynamics. The training is repeated multiple times until changes in the recurrent connectivity stabilize.

#### 2.1 Training synaptic drive

We modify the recurrent learning rule that was developed to train the recurrent connectivity of a network of rate units using Recursive Least Squares (RLS) algorithm^8,9^. Our learning rule extends the RLS algorithm to training the recurrent connectivity of spiking networks.

When learning the synaptic drive patterns, the objective is to find recurrent connectivity *W* that minimizes the cost function

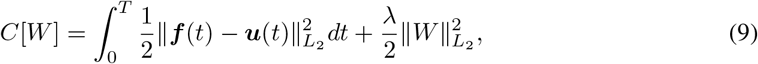

which measures the mean-square error between the targets and the synaptic drive over the time interval [0, *T*] plus a quadratic regularization term. To derive the learning rule, we use equation (8) to express *u* as a function of *W,* view the synaptic connections *W*_*i*1_,…, *W*_*iN*_ to neuron *i* to be the read-out weights that determine the synaptic drive *u*_*i*_, and apply the learning rule to the row vectors of *W*. To keep the recurrent connectivity sparse, learning occurs only on synaptic connections that are non-zero prior to training (see **Supplementary Notes** for details).

Let *w*_*i*_ (*t*) be the reduced row vector of *W*(*t*) consisting of elements that have non-zero connections to neuron *i* prior to training. Similarly, let *r*_*i*_(*t*) be a (column) vector of filtered spikes of presynaptic neurons that have non-zero connections to neuron *i*. The synaptic update to neuron *i* is

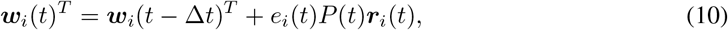

where the error term is

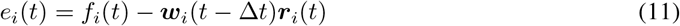

and the inverse of the correlation matrix of filtered spike trains is

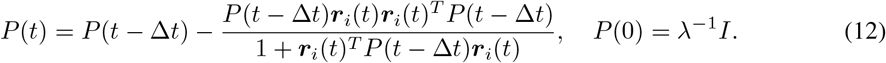

Finally, *W*(*t*) is obtained by concatenating the row vectors ***w****_i_*(*t*), *i* = 1,…, *N.*

#### 2.2 Training spiking rate

To train the spiking rate of neurons, we approximate the spike train *s*_*i*_(*t*) of neuron *i* with its spiking rate *φ*(*u*_*i*_(*t*) + *I*_*i*_) where *φ* is the current-to-rate transfer function of theta neuron model. For constant input,

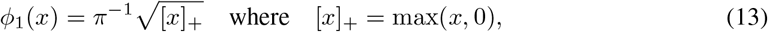

and for noisy input

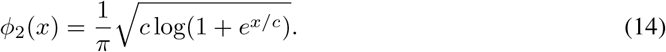

Since *φ*_2_ is a good approximation of *φ*_1_ and has a smooth transition around *x* = 0, we used *φ* ≡ *φ*_2_ with *c* = 0.1^58^. The objective is to find recurrent connectivity *W* that minimizes the cost function

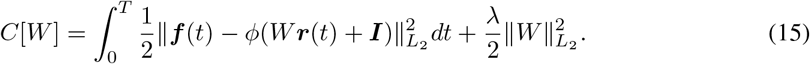

If we define *w*_*i*_ and *r*_*i*_ as before, we can derive the following synaptic update to neuron *i*

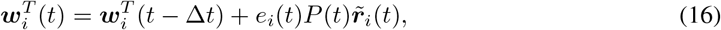

where the error term is

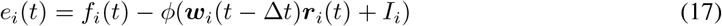

and

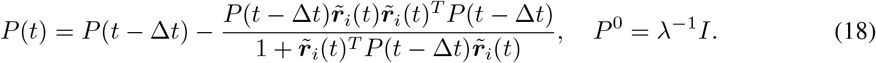

(see **Supplementary Notes** for details). Note that the nonlinear effects of the transfer function is included in

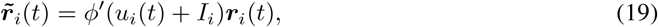

which scales the spiking activity of neuron *i* by its gain function *φ’*.

As before, *W*(*t*) is obtained by concatenating the row vectors *w*_*i*_(*t*), *i* = 1,…, *N*.

#### 3 Simulation parameters

Computer code. Example code is available at http://github.com/chrismkkim/SpikeLearning

**Figure 1**. A network of *N* = 200 neurons was connected randomly with probability *p* = 0.3 and the coupling strength was drawn from a Normal distribution with mean 0 and standard deviation 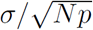 with *σ* = 4. In addition, the average of all non-zero synaptic connections to a neuron was subtracted from the connections to the neuron such that the summed coupling strength was precisely zero. Networks with balanced excitatory and inhibitory connections produced highly fluctuating synaptic and spiking activity in all neurons. The synaptic decay time was *τ*_*s*_ = 20 ms.

The target functions for the synaptic drive (**Fig. 1b**) were sine waves *f*(*t*) = *A* sin*(2π*(*t – T*_0_)/*T*_1_) where the amplitude *A,* initial phase *T*_0_, and period *T*_1_ were sampled uniformly from [0.5, 1.5], [0, 1000 ms] and [300 ms, 1000 ms], respectively. We generated *N* distinct target functions of length *T* = 1000 ms. The target functions for the spiking rate (**Fig. 1d**) were 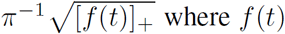 were the same synaptic drive patterns that have been generated.

Immediately before each training loop, every neuron was stimulated for 50 ms with constant external stimulus that had random amplitude sampled from [–1,1]. The same external stimulus was used across training loops. The recurrent connectivity was updated every ∆*t* = 2 ms during training using the learning rule derived from RLS algorithm and the learning rate was *λ* = 1. After training, the network was stimulated with the external stimulus to evoke the trained patterns. The performance was measured by calculating the average Pearson correlation between target functions and the evoked network response.

**Figure 2**. The initial network and target functions were generated as in **Figure 1** using the same parameters, but now the target functions consisted of two sets of *N* sine waves. To learn two sets of target patterns, the training loops alternated between two patterns, and immediately before each training loop, every neuron was stimulated for 50 ms with constant external stimuli that had random amplitudes, using a different stimulus for each pattern. Each target pattern was trained for 100 loops (i.e. total 200 training loops), synaptic update was every ∆*t* = 2 ms, and the learning rate was *λ* = 10. To evoke one of the target patterns after training, the network was stimulated with the external stimulus that was used to train that target pattern.

**Figure 3**. The network consisted of *N* = 500 neurons. The initial connectivity was sparsely connected with connection probability *p* = 0.3 and coupling strength was sampled from a Normal distribution with mean 0 and standard deviation 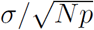 with σ = 1. The synaptic decay time was *τ*_*s*_ = 20 ms.

We considered three families of target functions with length *T* = 1000 ms. The complex periodic functions were defined as a product of two sine waves *f*(*t*) = *A* sin(2*π*(*t* – *T*_0_)/*T*_1_) sin(2*π*(*t* – *T*_0_)/*T*_2_) where *A, T*_0_, *T*_1_ and *T*_2_ were sampled randomly from intervals [0.5,1.5], [0,1000 ms], [500 ms, 1000 ms], and [100 ms, 500 ms], respectively. The chaotic rate activity was generated from a network of *N* randomly connected rate units, 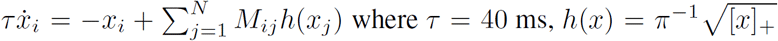 and *M*_*ij*_ is non-zero with probability *p* = 0.3 and is drawn from Gaussian distribution with mean zero and standard deviation 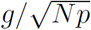 with *g* = 5. The Ornstein-Ulenbeck process was obtained by simulating, 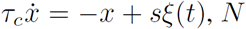 times with random initial conditions and different realizations of the white noise *ξ*(*t*) satisfying 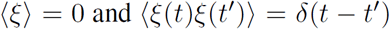. The decay time constant was *τ*_*c*_ = 200 ms, and the amplitude of target function was determined by *s* = 0.3.

The recurrent connectivity was updated every ∆*t* = 2 ms during training, the learning rate was *λ* = 1, and the training loop was repeated 30 times.

**Figure 4**. A balanced network had two populations where the excitatory population consisted of (1 – *f*) *N* neurons and the inhibitory population consisted of *f N* neurons with ratio *f* = 0.2 and network size *N* = 1000. Each neuron received *p*(1 – *f*)*N* excitatory connections with strength *J* and *pfN* inhibitory connections with strength –*gJ* from randomly selected excitatory and inhibitory neurons. The connection probability was set to *p* = 0.1 to have sparse connectivity. The relative strength of inhibition to excitation *g* was set to 5 so that the network was inhibition dominant^22^. In **Figure 4a-h**, the initial coupling strength *J* = 6 and synaptic decay time *τ*_*s*_ = 60 ms were adjusted to be large enough, so that the synaptic drive and spiking rate of individual neurons fluctuated strongly and slowly prior to training.

After running the initial network that started at random initial conditions for 3 seconds, we recorded the synaptic drive of all neurons for 2 seconds to harvest target trajectories that are innate to the balanced network. Then, the synaptic drive was trained to learn the innate trajectories, where synaptic update occurred every 10 ms, learning rate was *λ* = 10 and training loop was repeated 40 times. To respect Dale’s Law while training the network, we did not modify the synaptic connections if the synaptic update reversed the sign of original connections, either from excitatory to inhibitory or from inhibitory to excitatory. Moreover, the synaptic connections that attempted to change their signs were excluded in subsequent trainings. In **Figure 4h**, the initial and trained connectivity matrices were normalized by a factor 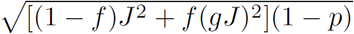 so that the spectral radius of the initial connectivity matrix is approximately 1, then we plotted the eigenvalue spectrum of the normalized matrices.

In **Figure 4i**, the coupling strength *J* was scanned from 1 to 6 in increments of 0.25, and the synaptic decay time *τ*_*s*_ was scanned from 5 ms to 100 ms in increments of 5 ms. To measure the accuracy of quasi-static approximation in untrained networks, we simulated the network dynamics for each pair of *J* and *τ*_*s*_, then calculated the average Person correlation between the predicted synaptic drive (equation (1)) and the actual synaptic drive. To measure the performance of trained networks, we repeated the training 10 times using different initial network configurations and innate trajectories, and calculated the Pearson correlation between the innate trajectories and the evoked network response for all 10 trainings. The heat map shows the best performance out of 10 trainings for each pair, *J* and *τ*_*s*_.

**Figure 5**. The initial connectivity was sparsely connected with connection probability *p* = 0.3 and the coupling strength was sampled from a Normal distribution with mean 0 and standard deviation 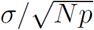 with *σ* = 1. The synaptic decay time was *τ*_*s*_ = 50 ms. There were in total *N* neurons in the network model, of which *N*_cor_ neurons, called cortical neurons, were trained to learn the spiking rate patterns of cortical neurons, and *N*_aux_ neurons, called auxiliary neurons, were trained to learn trajectories generated from OU process.

We used the trial-averaged spiking rates of neurons recorded in the anterior lateral motor cortex of mice engaged in motor planning and movement that lasted 4600 ms^5^. The data was available from the website CRCNS.ORG^40^. We selected *N*_cor_ = 227 neurons from the data set, whose average spiking rate during the behavioral task was greater than 5 Hz. Each cortical neuron in the network model was trained to learn the spiking rate pattern of one of the real cortical neurons.

To generate target rate functions for the auxiliary neurons, we simulated an OU process, 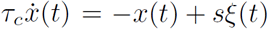, with *τ*_*c*_ = 800 ms and *s* = 0.1, then converted into spiking rate *φ*([*x*(*t*)]_+_) and low-pass filtered with decay time *τ*_*s*_ to make it smooth. Each auxiliary neuron was trained on 4600 ms-long target rate function that was generated with a random initial condition.

**Figures 6, 7**. Networks consisting of *N* = 500 neurons with no initial connections and synaptic decay time *τ*_*s*_ were trained to learn OU process with decay time *τ*_*c*_ and length *T*. In **Figure 6**, target length was fixed to *T* = 1000 ms while the time constants *τ*_*s*_ and *τ*_*c*_ were varied systematically from 10° ms to 5 • 10^2^ ms in log-scale. The trainings were repeated 5 times for each pair of *τ*_*s*_ and *τ*_*c*_ to find the average performance. In **Figure 7**, the synaptic decay time was fixed to *τ*_*s*_ = 20 ms and *T* was scanned from 250 ms to 5000 ms in increments of 250 ms, *τ*_*c*_ was scanned from 25 ms to 500 ms in increments of 25 ms, and *N* was scanned from 500 to 1000 in increments of 50.

To ensure that the network connectivity after training is sparse, synaptic learning occurred only on connections that were randomly selected with probability *p* = 0.3 prior to training. Recurrent connectivity was updated every ∆*t* = 2 ms during training, learning rate was λ = 1, and training loop was repeated 30 times. The average Pearson correlation between the target functions and the evoked synaptic activity was calculated to measure the network performance after training.

## Acknowledgments

This research was supported [in part] by the Intramural Research Program of the NIH, The National Institute of Diabetes and Digestive and Kidney Diseases (NIDDK).

## Supplementary notes

### 1 Training recurrent dynamics

Here, we derive the synaptic update rules for the synaptic drive and spiking rate trainings, (10) and (16). We use RLS algorithm^33^ to learn target functions *f*_*i*_(*t*), *i* = 1, 2, …, *N* defined on a time interval [0, *T*], and the synaptic update occurs at evenly spaced time points, 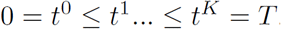.

In the following derivation, super-script *k* on a variable 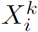 implies that *X* is evaluated at *t*^*k*^, and the sub-script *i* implies that *X* pertains to neuron *i*.

#### 1.1 Training synaptic drive

The cost function measures the discrepancy between the target functions *f*_*i*_(*t*) and the synaptic drive *u*_*i*_(*t*) for all *i* = 1, …, *N* at discrete time points *t*_0_, …, *t*_*K*_,

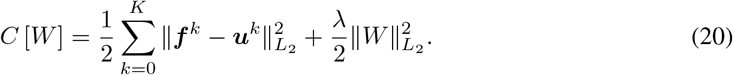

The Recursive Least Squares (RLS) algorithm solves the problem iteratively by finding a solution *W*^*n*^ to (20) at *t*^*n*^ and updating the solution at next time step *t*^*n*+1^. We do not directly find the entire matrix W^n^, but find each row of *W*^*n*^, i.e. synaptic connections to each neuron i that minimize the discrepancy between *u*_*i*_ and *f*_*i*_, then simply combine them to obtain *W*^*n*^.

To find the *i*^*th*^ row of *W*^*n*^, we denote it by 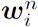 and rewrite the cost function for neuron *i* that evaluates the discrepancy between *f*_*i*_(*t*) and *u*_*i*_(*t*) on a time interval [0, *t*^*n*^],

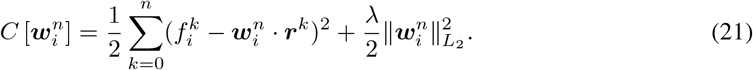

Calculating the gradient and setting it to 0, we obtain

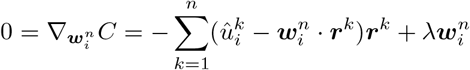

We express the equation concisely as follows.

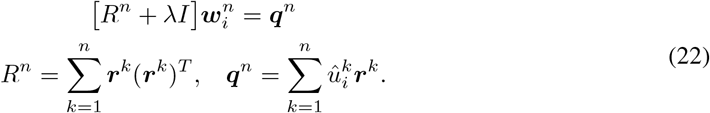

To find 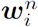 iteratively, we rewrite equation (22) up to *t*^*n*−1^,

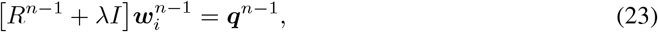

and subtract equations (22) and (23) to obtain

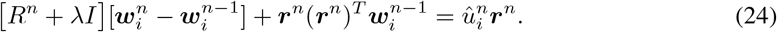

The update rule for 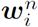 is then given by

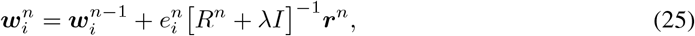

where the error term is

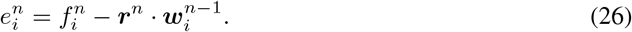

The matrix inverse 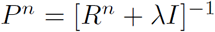 can be computed iteratively

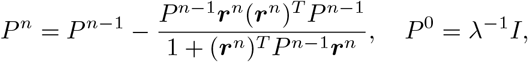

using the matrix identity

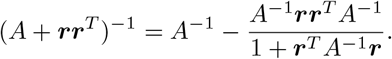

#### 1.2 Training spiking rate

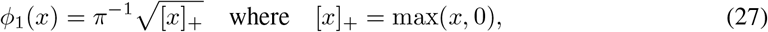

and for noisy input

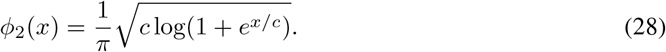

Since *φ*_2_ is a good approximation of *φ*_1_ and has a smooth transition around *x* = 0, we used *φ* ≡ *φ*_2_ with *c* = 0.1^58^.

If the synaptic update occurs at discrete time points, *t*^0^, …,*t*^*K*^, the objective is to find recurrent connectivity *W* that minimizes the cost function

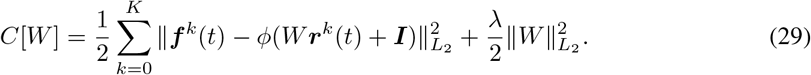

As in training the synaptic drive, we optimize the following cost function to train each row of *W*^*n*^ that evaluates the discrepancy between the spiking rate of neuron *i* and the target spiking rate *f*_*i*_ over a time interval [0, *t*_*n*_],

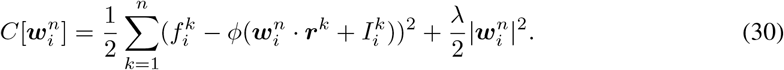

Calculating the gradient and setting it to zero, we obtain

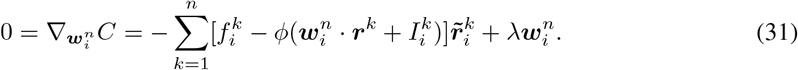

where

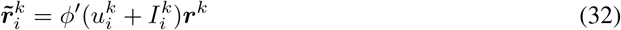

is the vector of filtered spike trains scaled by the gain of neuron *i*. Note that when evaluating *φ′* in equation (32), we use the approximation 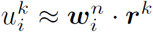 to avoid introducing nonlinear functions of 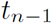.

To find an update rule for 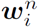, we rewrite equation (31) up to *t*_*n*–1_,

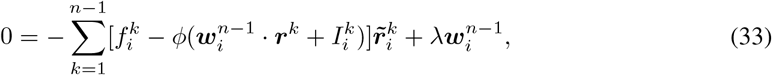

and subtract equations (31) and (33) and obtain

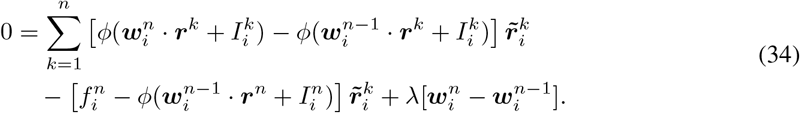

Since 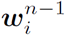 is updated by small increment, we can approximate the first line in equation (34),

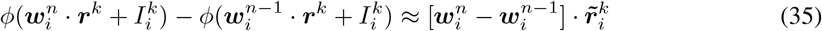

where we use the approximation 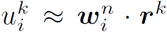 as before to evaluate the derivative 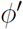. Substituting equation (35) to equation (34), we obtain the update rule

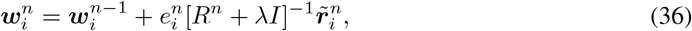

where the error is

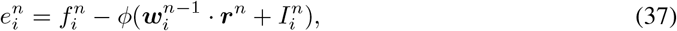

and the correlation matrix of the normalized spiking activity is

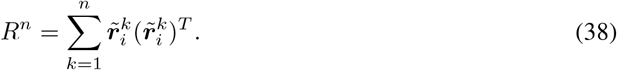

As shown above, the matrix inverse 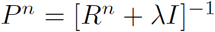 can be computed iteratively,

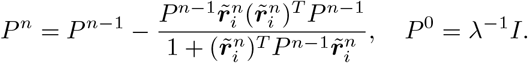

### 2 Mean field description of the quasi-static dynamics

We say that a network is in a quasi-static state if the synaptic drive to a neuron changes sufficiently slower than the dynamical time scale of neurons and synapses. Here, we use a formalism developed by Buice and Chow^41^ and derive equations (1) and (2), which provide a mean field description of the synaptic and spiking rate dynamics of neurons in the quasi-static state.

First, we recast single neuron dynamic equation (5) in terms of the empirical distribution of neuron’s phase η_i_(*θ, t*) = *δ*(*θ*_*i*_(*t*) – *θ*). Since the number of neurons in the network is conserved, we can write the Klimontovich equation for the phase distribution

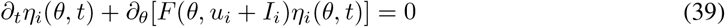

where *F*(*θ, I*) = 1 – cos *θ + I*(1 + cos *θ*). The synaptic drive equation (6) can be written in the form

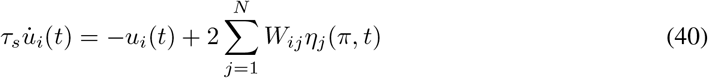

since 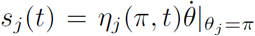 and 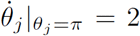for a theta neuron model. Equation (39), together with (40), fully describes the network dynamics.

Next, to obtain a mean field description of the spiking dynamics, we take the ensemble average prepared with different initial conditions and ignore the contribution of higher order moments resulting from nonlinear terms 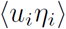. Then we obtain the mean field equation

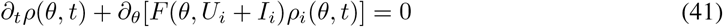

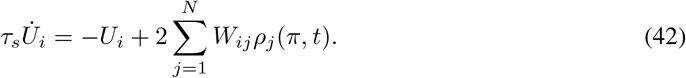

where 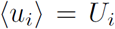 and 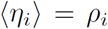. We note that the mean field equations (41) and (42) provide a good description of the trained network dynamics because *W* learns over repeatedly training trials and starting at random initial conditions, to minimize the error between target trajectories and actual neuron activity.

Now, we assume that the temporal dynamics of synaptic drive and neuron phase can be suppressed in the quasi-static state,

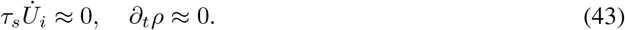

Substituting (43) to equation (41), but allowing *U*(*t*) to be time-dependent, we obtain the quasi-static solution of phase density

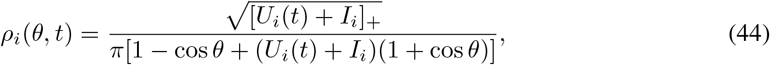

which has been normalized such that 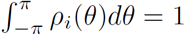, and the spiking rate of a neuron is given by

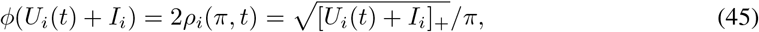

the current-to-rate transfer function of a theta neuron model. Substituting (43) and (45) to equation (42), we obtain a quasi-static solution of the synaptic drive

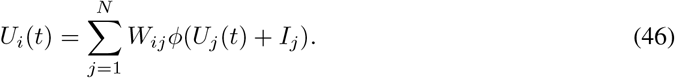

If we define the spiking rate of a neuron as *R*_*i*_(*t*) = *φ*(*U*_*i*_ + *I*_*i*_), we immediately obtain

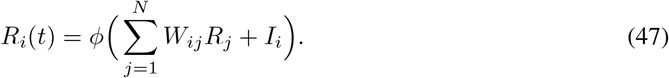

### 3 Analysis of learning error

In this section, we identify and analyze two types of learning errors, assuming that for sufficiently heterogeneous targets, (1) the learning rule finds a recurrent connectivity *W* that can generate target patterns if the quasi-static condition holds, and (2) the mean field description of the spiking network dynamics is accurate due to the error function and repeated training trials. These assumptions imply that equations (46) and (47) hold for the target patterns *U*_*i*_(*t*) and the trained *W*. We show that learning errors arise when our assumptions become inaccurate, hence the network dynamics described by equations (46) and (47) deviate from the actual spiking network dynamics. As we will see, tracking error is prevalent if the target is not an exact solution of the mean field dynamics (i.e. quasi-static approximation fails), and the sampling error dominates if the discrete spikes do not accurately represent continuous targets (i.e. mean field approximation fails).

Suppose we are trying to learn a target 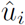 which obeys an Ornstein-Ulenbeck process

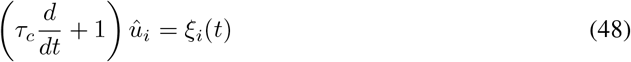

on a time interval 0 < *t < T* where *ξ*_*i*_(*t*) are independent white noise with zero mean and variance *σ*^2^. The time constant *τ*_*c*_ determines the temporal correlation of a target trajectory. In order for perfect training, the target dynamics (48) needs to be compatible with the network dynamics (6); in other words, there must exist a recurrent connectivity *W* such that the following equation

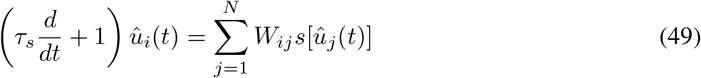

obtained by substituting the solution of (48) into (6) must hold for 0 < *t < T*. Here, 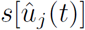 maps the synaptic drive 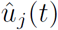 to the entire spike train *s*_*j*_ (*t*).

It is very difficult to find *W* that may solve equation (49) exactly since it requires fully understanding the solution space of a high dimensional system of nonlinear ordinary differential equations. Instead, we assume that the target patterns are quasi-static and the learning rule finds a recurrent connectivity *W* that satisfies

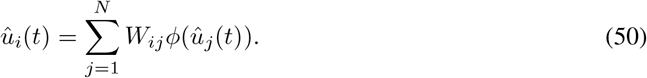

We then substitute equation (50) to equation (49) to estimate how the quasi-static mean field dynamics deviate from the actual spiking network dynamics. A straightforward calculation shows that

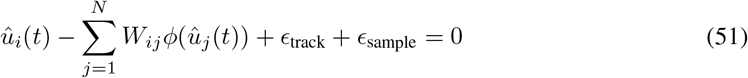

where we define the tracking and sampling errors as

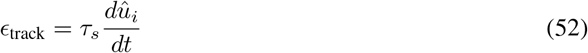

and

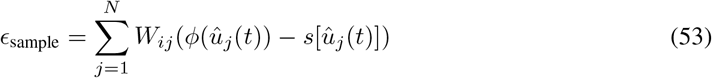

on the time interval 0 < *t < T*.

#### Tracking error

From its definition, *∊*_track_ captures the deviation of the quasi-static solution (50) from the exact solution of the mean field description obtained when *∊*_sample_ = 0. ∊_track_ becomes large if the quasi-static condition (43) fails and, in such network state, the synaptic dynamic is not able to “track” the target patterns, thus learning is obstructed. In the following, we estimate *∊*_track_ in terms of two time scales *τ*_*s*_ and *τ*_*c*_.

First, we take the Fourier transform of equation (52) and obtain

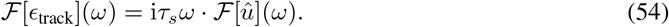

Next, normalize 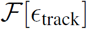 with respect to 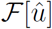 to estimate the tracking error for target patterns with different amplitudes, then compute the power of normalized tracking error.

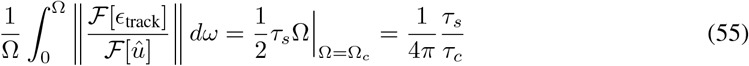

where 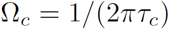 is the cut-off frequency of the power spectrum of a Gaussian process, *S*_*GP*_ (*ω*) = 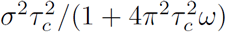. Thus, the tracking error scales with *τ*_*s*_/*τ*_c_.

#### Sampling error

*∊*_sample_ captures how the actual representation of target patterns in terms of spikes deviates from their continuous representation in terms of rate functions. In the following, we estimate *∊*_sample_ in terms of *τ*_*s*_ and *N* under the assumption that the continuous representation provides an accurate description of the target patterns.

We low-pass filtered ∊_sample_ to estimate the sampling error since the synaptic drive (i.e. the target variable in this estimate) is a *W* weighted sum of filtered spikes with width that scales with *τ*_*s*_. If the spike trains of neurons are uncorrelated (i.e. cross product terms are negligible),

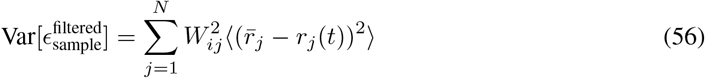

where *r*_*j*_(*t*) is the filtered spike train and 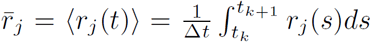 is the empirical estimate of mean spiking rate on a short time interval.

First, we calculate the fluctuation of filtered spike trains under the assumption that a neuron generates spikes sparsely, hence the filtered spikes are non-overlapping. Let 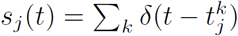 be a spike train of neuron *j* and the filtered spike train 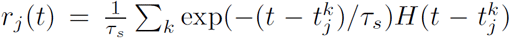. Then, the rate fluctuation of neuron *j* is

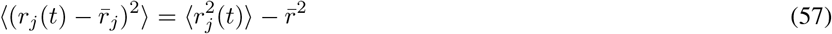

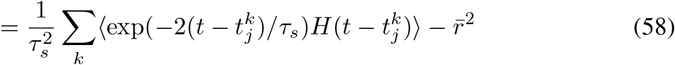

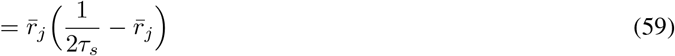

where *k* is summed over the average number of spikes, 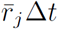, generated in the time interval of length ∆*t*.

Next, to estimate the effect of network size on the sampling error, we examined equation (50) and observed that *O* (*W*) ~ 1/*N*. This follows from that, for pre-determined target patterns, *O*(*U*), *O*(φ(*U*)) ~ 1 regardless of the network size, hence *O*(*W*) must scale with 1/*N* in order for both sides of the equation to be compatible. If the network is dense, i.e. the number of synaptic connections to a neuron is *pN* on average, then the sampling error scales as follows.

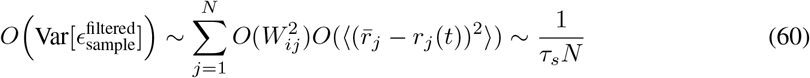

### Supplementary figures

**Figure S1:**
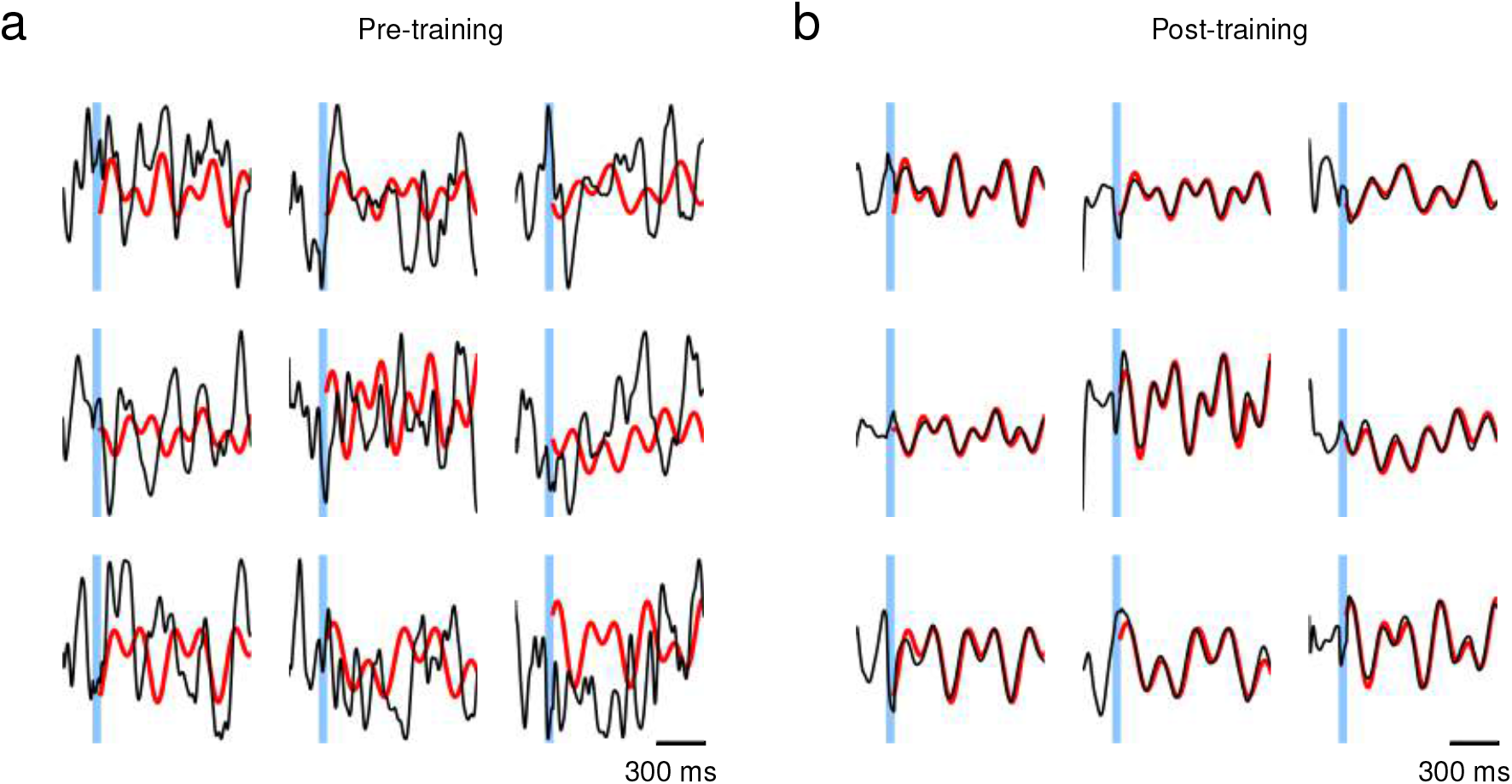
Learning arbitrarily complex target patterns in a network of rate-based neurons. The network dynamics obey 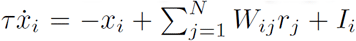 where 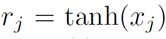. The synaptic current *x*_*i*_ to every neuron in the network was trained to follow complex periodic functions *f*(*t*) = *A* sin(*2*π(*t – T*_0_)*/T*_1_) sin(2π(*t* – *T*_0_)/*T*_2_) where the initial phase *T*_0_ and frequencies *T*_1_, *T*_2_ were selected randomly. The elements of initial connectivity matrix *W*_*ij*_ were drawn from a Gaussian distribution with mean zero and standard deviation 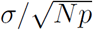 where *σ* = 2 was strong enough to induce chaotic dynamics; Network size *N* = 500, connection probability between neurons *p* = 0.3, and time constant *τ* = 10 ms. External input *I*_*i*_ with constant random amplitude was applied to each neuron for 50 ms (blue) and was set to zero elsewhere. (a) Before training, the network is in chaotic regime and the synaptic current (black) of individual neurons fluctuates irregularly. (b) After learning to follow the target trajectories (red), the synaptic current tracks the target pattern closely in response to the external stimulus.

**Figure S2:**
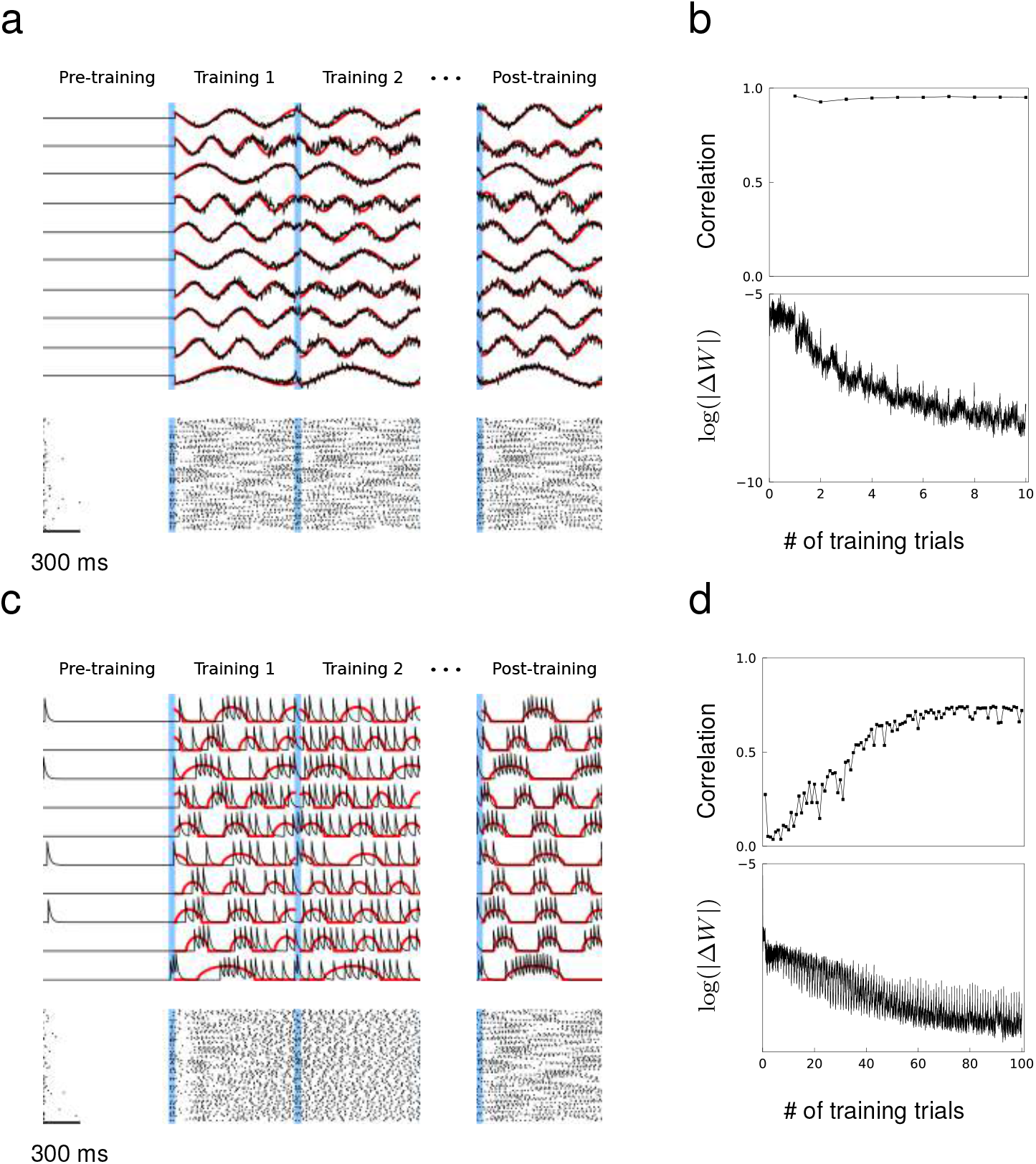
Training a network that has no initial connections. The coupling strength of the initial recurrent connectivity is zero, and, prior to training, no synaptic or spiking activity appears beyond the first few hundred milliseconds. (a) Training synaptic drive patterns using the RLS algorithm. Black curves show the actual synaptic drive of 10 neurons and red curves show the target outputs. Blue shows the 100 ms external stimulus. (b) Correlation between synaptic drive and target function (top) and the Frobenius norm of changes in recurrent connectivity normalized to initial connectivity during training (botom). (c)-(d) Same as in (a) and (b), but spiking rate patterns are trained.

**Figure S3:**
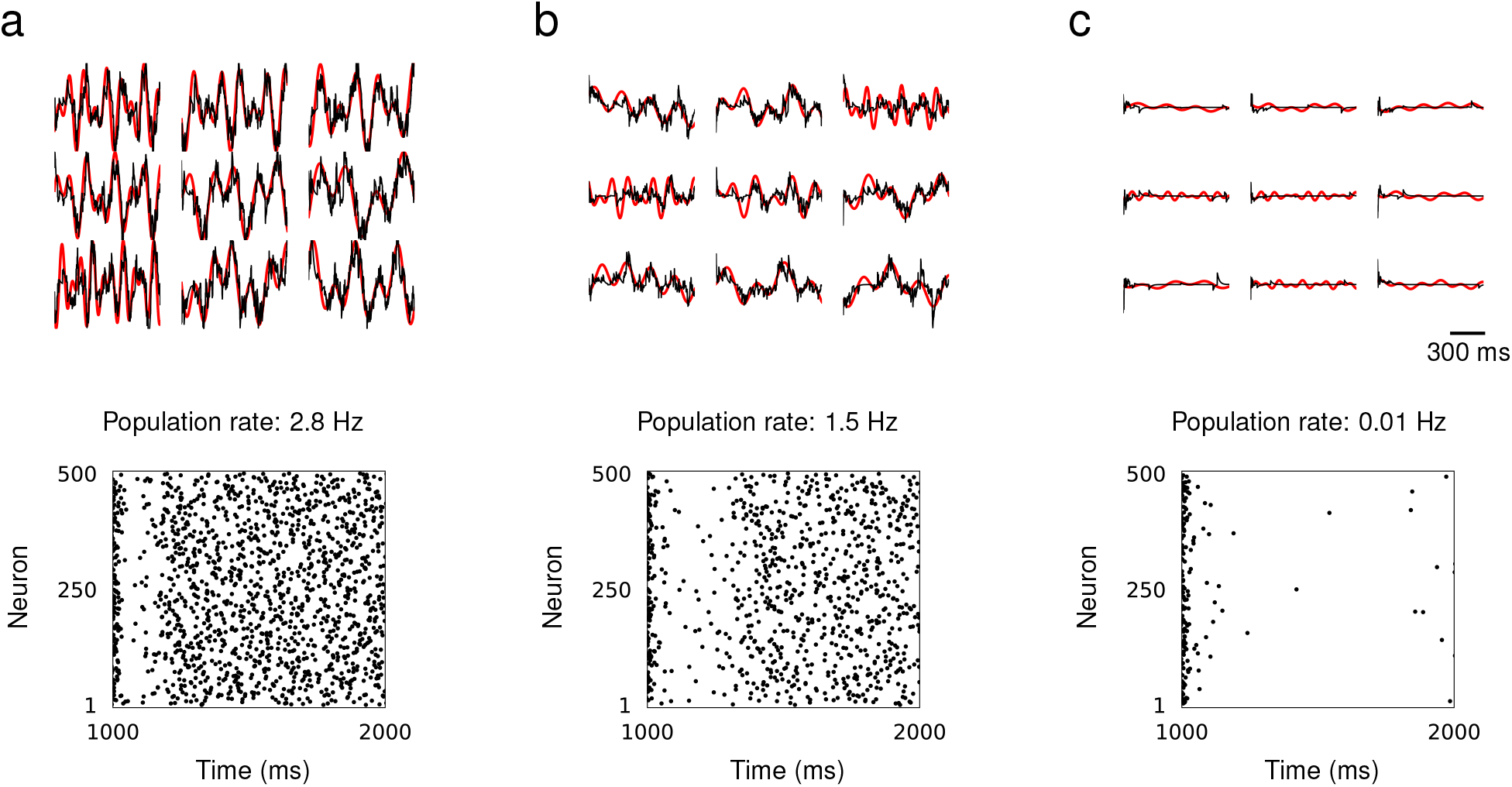
Learning target patterns with low population spiking rate. The synaptic drive of networks consisting of 500 neurons were trained to learn complex periodic functions *f*(*t*) = *A* sin(*2*π(*t – T*_0_)/*T*_1_) sin(2π(*t* – *T*_0_)/*T*_2_) where the initial phase *T*_0_ and frequencies *T*_1_, *T*2 were selected randomly from [500 ms, 1000 ms]. (a) The amplitude *A* = 0.1, resulting in population spiking rate 2.8 Hz in trained window. (b) The amplitude *A* = 0.05, resulting in population spiking rate 1.5 Hz in trained window. (c) The amplitude *A* = 0.01, resulting in population spiking rate 0.01 Hz in trained window and learning fails.

**Figure S4:**
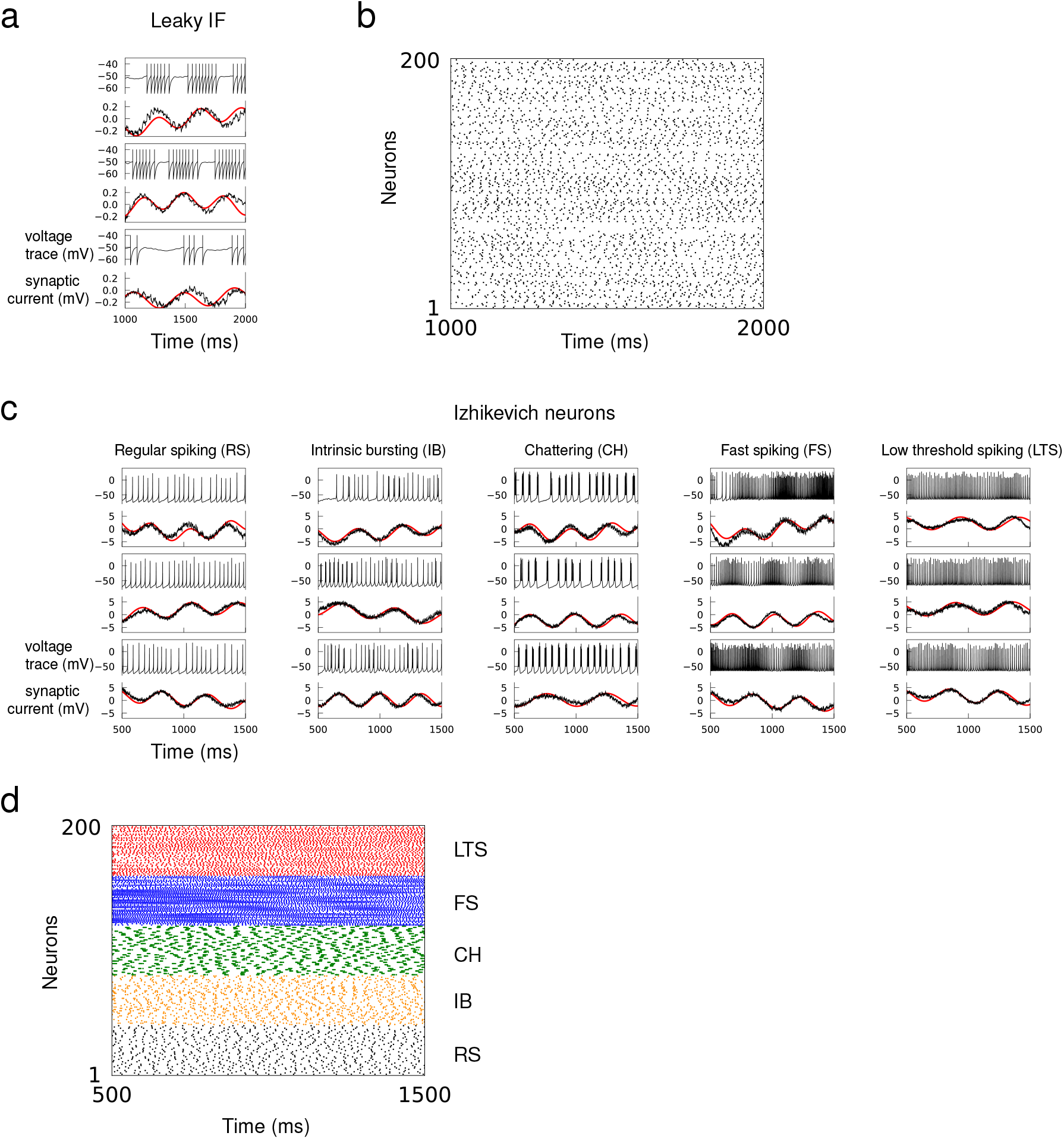
Learning recurrent dynamics with leaky integrate-and-fire and Izhikevich neuron models. Synaptic drive of a network of spiking neurons were trained to follow 1000 ms long targets 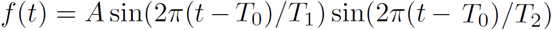 where *T*_0_, *T*_1_ and *T*_2_ were selected uniformly from the interval [500 ms, 1000 ms]. (**a**) Network consisted of *N* = 200 leaky integrate-and-fire neuron models, whose membrane potential obeys 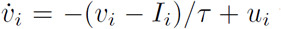 with a time constant *τ* = 10 ms; the neuron spikes when *v*_*i*_ exceeds spike threshold *v*_*thr*_ = –50 mV then *v*_*i*_ is reset to *v*_*res*_ = –65 mV. Red curves show the target pattern and black curves show the voltage trace and synaptic drive of a trained network. (b) Spike rastergram of a trained leaky integrate-and-fire neuron network generating the synaptic drive patterns. (c) Network consisted of *N* = 200 Izhikevich neurons, whose dynamics are described by two equations 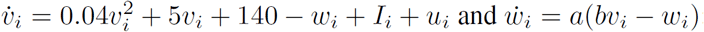; the neuron spikes when *v*_*i*_ exceeds 30 mV, then *v*_*i*_ is reset to *c* and *w*_*i*_ is reset to *w*_*i*_ + *d*. Neuron parameters *a, b, c* and *d* were selected as in the original study^59^ so that there were equal numbers of regular spiking, intrinsic bursting, chattering, fast spiking and low threshold spiking neurons. Synaptic current *u*_*i*_ is modeled as in equations (6) for all neuron models with synaptic decay time *τ*_*s*_ = 30 ms. Red curves show the target patterns and black curves show the voltage trace and synaptic drive of a trained network. (**d**) Spike rastergram of a trained Izhikevich neuron network showing the trained response of different cell types.

**Figure S5:**
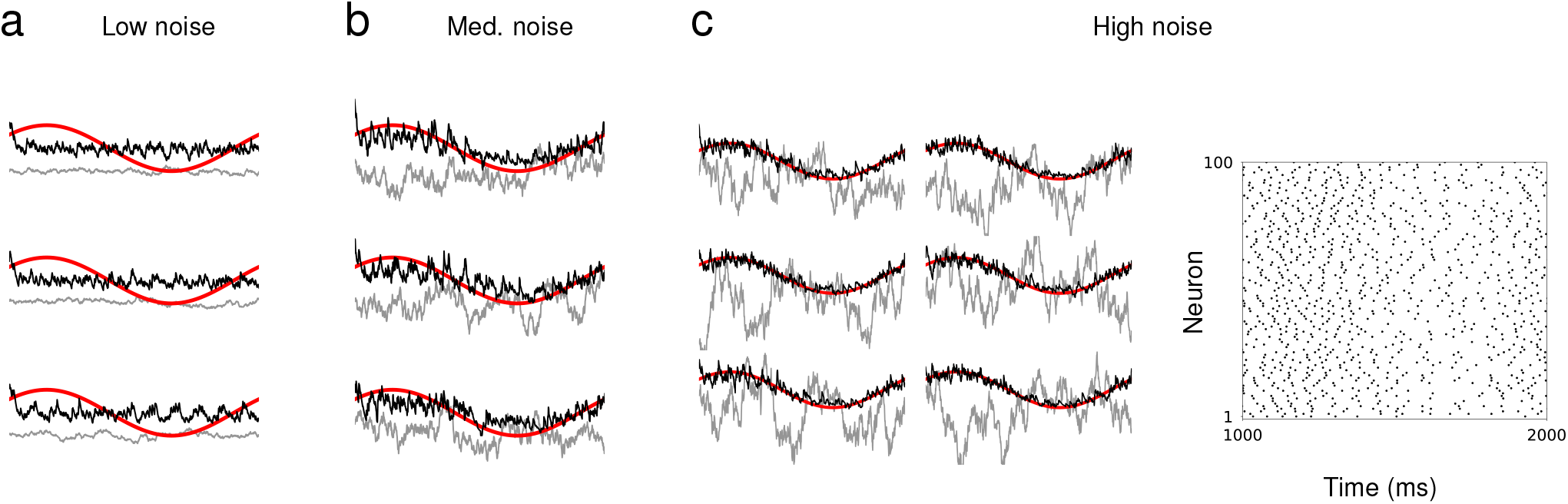
Synaptic drive of a network of neurons is trained to learn an identical sine wave while external noise generated independently from OU process is injected to individual neurons. The same external noise (gray curves) is applied repeatedly during and after training. **(a)-(b)** The amplitude of external noise is varied from (a) low, (b) medium to (c) high. The target sine wave is shown in red and the synaptic drive of neurons are shown in black. The raster plot in (c) shows the ensemble of spike trains of a successfully trained network with strong external noise.

